# Evaluating metagenomic analyses for undercharacterized environments: what’s needed to light up the microbial dark matter?

**DOI:** 10.1101/2024.11.08.622677

**Authors:** William A. Nickols, Lauren J. McIver, Aaron Walsh, Yancong Zhang, Jacob T. Nearing, Francesco Asnicar, Michal Punčochář, Nicola Segata, Long H. Nguyen, Erica M. Hartmann, Eric A. Franzosa, Curtis Huttenhower, Kelsey N. Thompson

## Abstract

Non-human-associated microbial communities play important biological roles, but they remain less understood than human-associated communities. Here, we assess the impact of key environmental sample properties on a variety of state-of-the-art metagenomic analysis methods. In simulated datasets, all methods performed similarly at high taxonomic ranks, but newer marker-based methods incorporating metagenomic assembled genomes outperformed others at lower taxonomic levels. In real environmental data, taxonomic profiles assigned to the same sample by different methods showed little agreement at lower taxonomic levels, but the methods agreed better on community diversity estimates and estimates of the relationships between environmental parameters and microbial profiles.

## Background

Microbes contribute many essential functions to their ecosystems, from rivers and oceans to humans and other mammalian hosts(1–5). Advances in sequencing over the last few decades have driven a rapid increase in our understanding of these essential microbes and their role in ecosystems(6–8). However, historically, a majority of microbiome-focused work has been directed at human-associated communities, resulting in a knowledge and software bias towards those communities and their microbes. For instance, a recent publication found that 75% of non-human host-associated genomes in gut microbiome samples had never been previously described(4), representing considerable “microbial dark matter”(9). Likewise, another recent publication found that 83% of free-living polar ocean metagenomic assembled genomes (MAGs) were unclassified at the species level(10), and a study of Baltic sea samples reported that 91% (320/352) of identified species had not been previously characterized(11). Continually advancing taxonomic profiling methods are necessary to close this knowledge gap, and evaluations of these methods in highly uncharacterized settings are necessary to inform future studies.

Two broad strategies exist for profiling microbial communities: reference and assembly-based methods. Reference-based methods can reliably profile communities with shallower sequence coverage (approximately 0.05X)(12), but they inherently miss novel taxa that lack references(13). Conversely, assembly methods can construct genomes for novel species, enabling further characterization, but they often require 10x sequence coverage or more(14). This biases profiling to highly abundant microbes (e.g. microbes with greater than ∼3% relative abundance in a 10 million read library assuming a 4 million base pair genome and 150 base reads). Because non-human associated communities are often enriched in novel uncultured microbes (those lacking an isolate in a reference database), environmental research has often relied heavily on assembly-based methods to profile communities(4, 7, 8). To combat the need for such deep sequencing, studies often rely on pooled assembly of multiple samples (15, 16), but this can reduce our understanding of natural variation(17, 18) and can still miss sufficiently rare or hard-to-assemble microbes. Additionally, as these assemblies are incorporated into reference databases, reference methods can become biased towards highly abundant microbes. While recent benchmarking analyses have included performance in under-characterized communities(19–21), no prior work has specifically evaluated the optimal strategy for profiling under-characterized microbial communities.

Here, we seek to address this gap by comparing the performance of state-of-the-art reference-and assembly-based profiling methods on realistic synthetic environmental communities and real environmental (non-human associated) samples. To this end, we evaluated the methods on 300 synthetic taxonomic profiles over a range of sample parameters (e.g., sequencing depth, proportion of uncharacterized species, and environment) and on 348 real environmental samples from distinct ecosystem types. The synthetic evaluation revealed that assembly-based methods suffered in communities that were complex or shallowly sequenced, and the performance of reference-based methods deteriorated at higher, still-characterized taxonomic levels when more of the underlying species were uncharacterized. The evaluation on real samples showed significant variation in taxonomic profiles assigned to the same sample but moderate agreement on downstream metrics of community structure and taxa-metadata associations. Ultimately, this analysis will help inform the interpretation of environmental microbial communities and guide future development of taxonomic profilers, especially as they focus more on environmental communities.

## Results

### Diverse environmental datasets were curated for realistic comparisons

To assess the performance of state-of-the-art taxonomic profilers in non-human environmental communities, we tested the profilers on both synthetic environmental and real data. For the synthetic communities, we curated a set of characterized and uncharacterized genomes, constructed abundance profiles based on realistic environmental sample parameters (e.g. number of species), and generated synthetic short reads with CAMISIM(22) (**Fig. 1A-C; Methods**).

**Figure 1:**
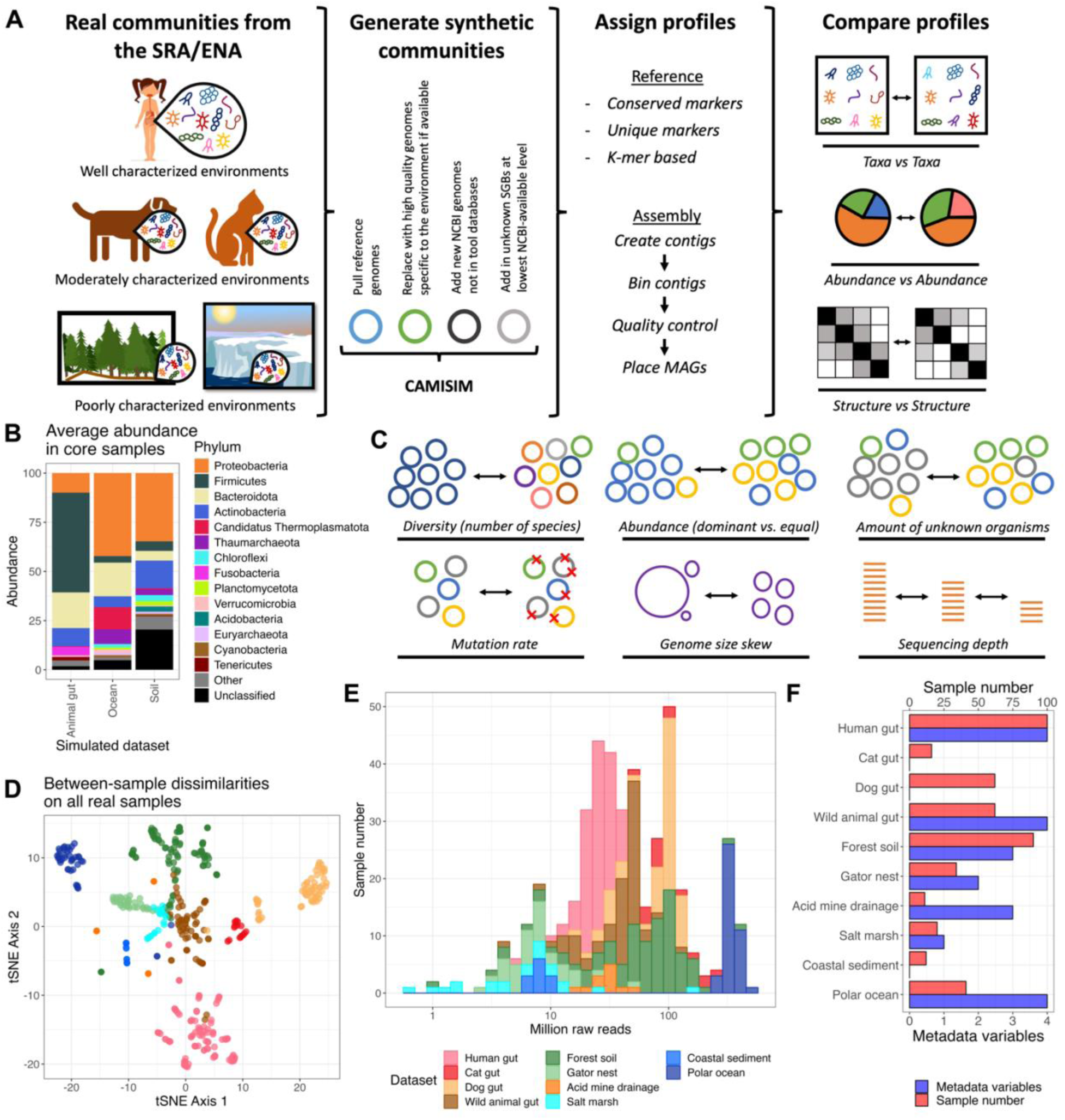
Workflow and dataset overview. **A.** Publicly available sequencing projects were evaluated with a variety of state-of-the-art metagenomic methods, and their resulting profiles were compared. In addition, these real samples informed synthetic read generation, and these synthetic profiles were also evaluated. **B.** Synthetic samples represented real microbial communities in their taxa and composition. Average abundance per phylum in each simulated dataset for core samples most representative of the environment. **C.** Relevant community and sample parameters were varied between samples in each simulated dataset. **D.** Datasets differed in underlying taxonomic composition. tSNE ordination of between-sample dissimilarities on real data determined by Simka. **E.** Samples spanned a wide range of sequencing depths. Read count per sample. **F.** Datasets differed in how much metadata and how many samples were available. Per dataset, the number of samples available and the number of metadata variables publicly available and included for evaluation.

To represent diverse environments, we curated 348 real environmental samples from various sources including high-diversity soil, moderate-diversity water, low-diversity acid mine drainage, and moderately characterized animal gut environments (**Fig. 1A**, **1F** and **Supplemental Table 1**)(23–30). We compared the performance of taxonomic profiling methods on this diverse array of samples to performance on 100 human gut samples from participants without Inflammatory Bowel Disease in the Human Microbiome Project 2 (HMP2)(31). Together, these datasets represented environments with different levels of characterization: the human gut samples represent well-characterized environments (average 41% uncharacterized abundance across human samples assigned by reference methods at the species level); the animal gut samples represent moderately characterized environments (cat gut 48%, dog gut 64%, wild animal gut 70%); and the purely environmental samples represent poorly characterized environments (acid mine drainage 91%, coastal sediment 91%, forest soil 93%, gator nest soil 82%, salt marsh 89%, polar ocean 78%). Using the reference-independent method Simka(32), we ensured that the evaluated communities varied considerably in underlying taxonomic composition (**Fig. 1D**). Additionally, these datasets differed in their sequencing depths (**Fig. 1E**) and metadata availability (for use in association testing; **Fig. 1F**). Still, all samples were sequenced on Illumina short-read instruments, in most cases using paired-end sequencing (**Supplemental Table 1**).

For this evaluation, we focused on taxonomic assignment methods that were frequently cited and used in metagenomic literature (**Supplemental Table 2**). The reference-based methods included were Centrifuge(33) and Kraken 2/Bracken 2(34, 35) as *k*-mer-based methods, mOTUs3(19) and Metaxa2(36) as universal marker gene methods, and MetaPhlAn 2/3/4(20, 37, 38) as the unique marker method (**Methods**). For assembly-based methods, we used two very similar pipelines with contig construction by either MEGAHIT(39) or metaSPAdes(40), genome binning by MetaBAT(41), binning quality control by CheckM2 (42), and taxonomic assignment of the metagenomic assembled genome bins (MAGs) by PhyloPhlAn(43) or GTDB-Tk(44) (**Methods**). To avoid biasing the results towards any particular method, no parameters were tuned for any method, and the most recent default databases (as of June 2022) were downloaded or constructed for each method (**Methods**). Even after harmonizing taxonomic naming (**Methods**), the databases still differed significantly in which taxa they contained and how many taxa overlapped with other databases (**Supplemental Fig. 1**). While some previous evaluations have used standardized databases to avoid these differences(45), we sought to evaluate each taxonomic classification strategy in the way it would be used practically and therefore considered the standard database to be an integral part of the method.

### Taxonomic profiling methods differ in accuracy on simulated environmental profiles

To assess the accuracy of the taxonomic profiling methods, we generated synthetic reads from known compositions of microbes representative of ocean, soil, and animal gut community structures; profiled each synthetic sample with each taxonomic assignment method; and compared the assigned profiles to the true profiles (**Methods**). For the first set of analyses, we focused on “core samples” constructed with the most realistic sample parameters: 300 species, 7.5 giga-base pairs of sequencing (50 million 150 bp reads), 75% unknown species for soil and ocean samples and 50% unknown species for animal gut samples, and an underlying log normal distribution for abundances (parameters µ=-3 and σ=1) converted to relative abundances. In order to model the high proportion of uncharacterized species in non-human samples (**Supplemental Fig. 2**), we constructed the default samples so that 50% (ocean and animal gut) or 75% (soil) of their abundance was from species not represented in NCBI (**Methods** and **Fig. 2A**). The methods’ default databases often included only a subset of the species available in NCBI, but in the simulated profiles, the abundance of taxa that were available in NCBI but not included in the profilers’ databases was usually small and mostly depended on the database’s age (**Supplemental Fig. 3**). In core samples, such taxa contributed at most 35.7% abundance (mean 8.3%) for any method at any taxonomic level below the kingdom level. To compare the methods impartially, our analyses focused on only these “NCBI-available” taxa (**Methods**).

**Figure 2:**
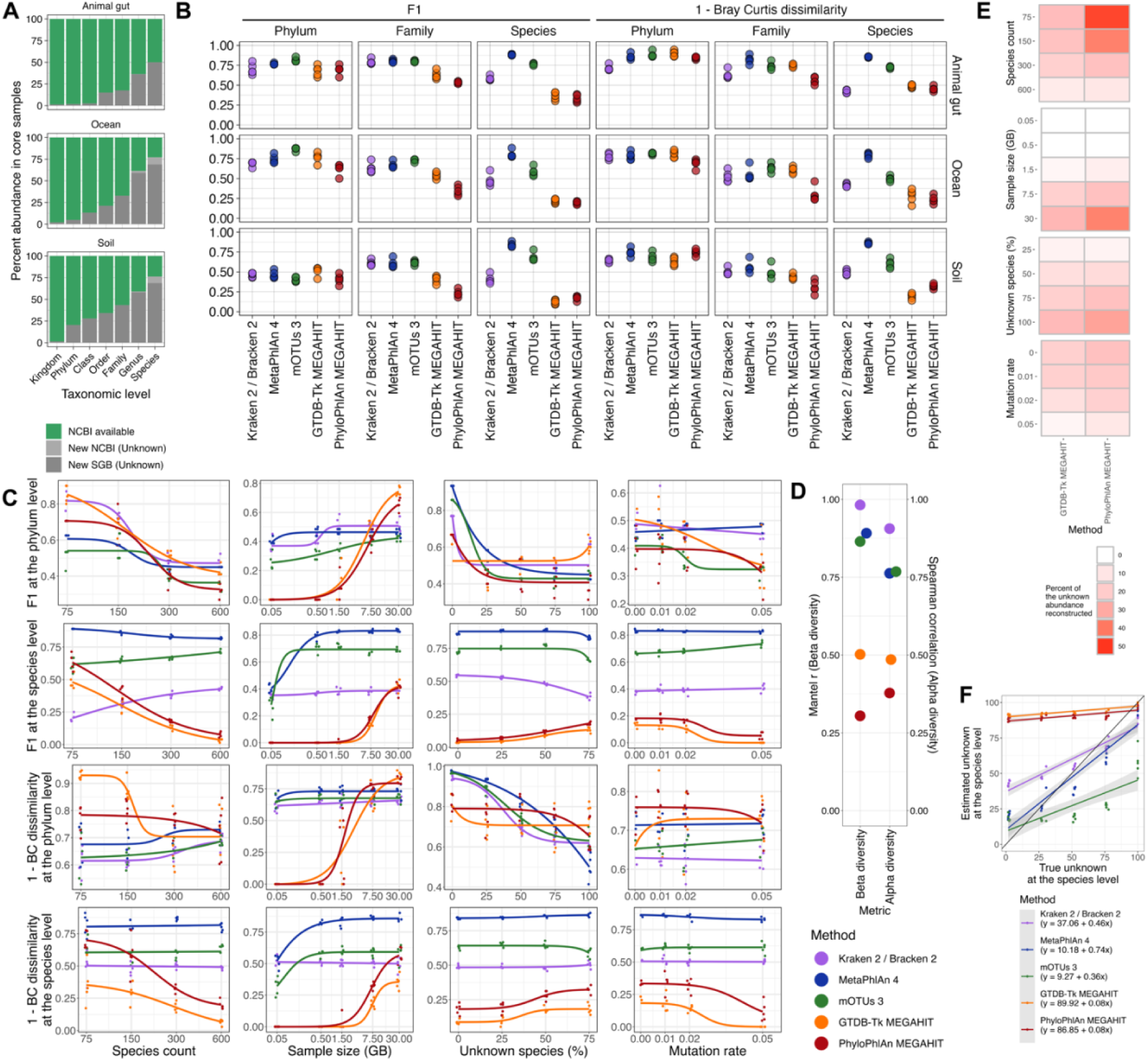
Taxonomic profiling methods differ in accuracy on simulated profiles. **A.** In core samples, the abundance of taxa with historically available NCBI identifiers (NCBI available), NCBI identifiers added since the most recent method database update (new NCBI), or no NCBI identifier (new SGB). F1 and Bray-Curtis dissimilarity scores were computed by comparing assigned profiles only to the NCBI available segment. **B.** Methods differed in accuracy on core samples most representative of each environment. F1 and Bray-Curtis dissimilarity relative to the true profile for core samples by simulated dataset. **C.** Method accuracy differed with sample parameters. F1 and Bray-Curtis dissimilarity by sample parameter for the simulated soil dataset (5 replicates per parameter setting). Only one parameter at a time was changed from the core value. 4 parameter logistic curves were fit for each. Panels **C-F** show only the simulated soil dataset, but similar trends emerged in the ocean and animal gut samples (**Supplemental Fig. 4-5**). **D.** Method accuracy differed on community structure metrics. (Left) Mantel test of the true species BC dissimilarity matrix against the reconstructed BC dissimilarity matrices. (Right) Spearman correlation between true and reconstructed inverse Simpson indices at the species level. For both metrics, all samples in the simulated soil dataset were included with their full profiles (NCBI available and not). **E.** Sample parameters affected assembly methods’ abilities to reconstruct uncharacterized genomes. Percent abundance of species with genomes lacking species level characterization successfully reconstructed and assigned a proper taxonomy. **F.** Methods differed in their abilities to predict the abundance of uncharacterized taxa. Estimated proportion of the abundance unknown at the species level versus the true abundance of species new to NCBI or from new SGBs.

We first evaluated the ability of the profilers to assign the correct microbes to a sample (using the F1 between the true and assigned profiles) and estimate the correct microbial abundances (using Bray-Curtis (BC) dissimilarity). Compared to the unweighted F1 score, the Bray-Curtis dissimilarity helps identify when the majority of a sample’s abundance is correctly identified even if low-abundance species are misidentified. On the core samples, all methods performed similarly at the phylum level for both metrics, but this accuracy differed considerably by environment (average F1 0.72 (animal gut), 0.69 (ocean), 0.43 (soil); average BC dissimilarity 0.18 (animal gut), 0.24 (ocean), 0.33 (soil); **Fig. 2B**). At the family level, reference-based methods produced more accurate profiles than assembly-based methods, particularly on the F1 metric (average F1 across all datasets 0.63 versus 0.42 respectively), an effect mostly attributable to the lower recall of assembly-based methods. At the species level, MetaPhlAn 4 exhibited the highest accuracy on both the F1 and BC dissimilarity metrics across all environments (average F1 0.84; BC dissimilarity 0.16) with both high precision and high recall. Modest accuracy was also achieved by the profiler mOTUs3 (F1 0.68; BC dissimilarity 0.39) and to a lesser extent Kraken 2/Bracken 2 (F1 0.49; BC dissimilarity 0.56). The F1 result for mOTUs3 was driven by its high precision but lower recall, while the F1 for Kraken 2/Bracken 2 came from both moderate recall and precision (**Supplemental Fig. 4-6**). The higher accuracy and recall of MetaPhlAn 4 and mOTUs3 is likely due to their inclusion of environmentally-sourced MAGs in their databases, these being the only reference methods that did not rely on isolate references. Assembly-based methods performed poorly on both metrics due to their very low recall (range of F1 averages 0.16-0.24; BC dissimilarity 0.66-0.73). This is consistent with the fact that many things must go right for an assembly method to perform well: reads must be assembled successfully, contigs must be binned correctly, bins must be taxonomically placed accurately, and per-MAG abundance must be estimated well.

To assess how naturally varying features of a sample affect the profiling accuracy, we varied the underlying species diversity, sequencing depth, proportion of uncharacterized species, mutation rate, genome size skew, and abundance skew in each environment (**Methods**). For samples with more species, assembly-based methods exhibited decreased recall at both the phylum and species levels because reconstructing MAGs for all species became more difficult as individual genome coverage decreased (**Fig. 2C**, **Supplemental Fig. 4-6**). Sequencing depth did not affect reference-based methods when samples had at least 0.5 giga-base pairs, though it affected all methods except *k*-mer-based methods below 0.5 giga-base pairs (**Fig. 2C**). Like species richness, sequencing depth was important for assembly-based methods since they required sufficient genome coverage to construct medium and high quality genomes: samples with fewer than 7.5 giga-base pairs resulted in very low F1 scores and high BC dissimilarity from the true profile. Samples with more extreme abundance skews yielded lower recall for all methods as more taxa contributed few reads (**Supplemental Fig. 4-6**). However, more skewed abundances produced lower BC dissimilarity for assembly methods as the genomes that could best be reconstructed accounted for more of the abundance. Including larger proportions of uncharacterized species changed species-level accuracy very little in this analysis since we restricted the evaluation to NCBI-available taxa (**Fig. 2C**). However, at the phylum level, accuracy decreased when more species were unknown even though those species’ upper-level taxonomies were still known. That is, the methods were often unable to accurately assign higher-level taxonomies to species that were not in their databases even though the higher-level taxonomic groups *were* in their databases. The impact of larger unknown proportions on assembly-based methods was highly variable depending on the environment, likely due to database biases (**Supplemental Fig. 4-6**). The random point mutation rate generally had a negligible effect on the methods’ accuracies at least within the range of mutation rates evaluated (**Fig. 2C**). We also sought to determine the effect of genome size variability on accuracy because a recent study reiterated that read classifiers (like Kraken) cannot be compared directly to abundance classifiers (like MetaPhlAn) because of genome size differences(46). However, we found that the variability in genome sizes actually had little impact on the BC dissimilarity of read classifiers compared to abundance classifiers (**Supplemental Fig. 4-6**). This could be due to the fact that, unlike in the previous study, the profiles assigned by most methods here were already so different from the true profiles that additional inaccuracies from genome size variation had little effect.

Beyond taxonomic profile accuracy, a method could remain useful if it accurately reconstructs community structure metrics such as alpha or beta diversity. To evaluate the methods’ performances on these tasks, we computed the species pairwise BC dissimilarity matrices for the true and assigned profiles. In all datasets, the *k*-mer methods with a 0.05% abundance filter most accurately reconstructed the true BC dissimilarity matrices (Mantel r 0.95-0.99 across *k*-mer methods and across datasets); the other reference-based methods except for Metaxa2 constructed nearly as accurate matrices (Mantel r 0.79-0.94); and the assembly-based methods and Metaxa2 constructed the least accurate matrices (Mantel r 0.22-0.71) (**Fig. 2D** and **Supplemental Fig. 7**). We then computed the inverse Simpson alpha diversity for each true and assigned profile. In soil and animal gut samples, the *k*-mer-based methods produced the most accurate alpha diversity estimates (Spearman correlation 0.79–0.94 across *k*-mer-based methods and across datasets); the other reference-based methods produced slightly less accurate estimates (Spearman correlation 0.59-0.77); and the assembly-based methods produced the least accurate estimates (Spearman correlation 0.34-0.57) (**Fig. 2D** and **Supplemental Fig. 7**). In the ocean samples, Metaxa2 produced the most accurate alpha diversity estimate (Spearman correlation 0.85), followed by the other reference-based methods (Spearman correlation 0.71-0.74) and the assembly-based methods (0.33-0.36) (**Supplemental Fig. 7**). These results suggest that *k*-mer methods, closely followed by the other reference based-methods, best capture structural patterns in uncharacterized microbial communities.

Despite the lower performance of assembly-based strategies thus far, they are the only methods that can assess novel taxa, an essential task in most environmental analyses. Therefore, we evaluated how sample parameters impacted the proportion of uncharacterized species the assembly-based methods successfully reconstructed (**Methods**). As expected, the main determinant of genome reconstruction was genome coverage: a larger abundance of uncharacterized genomes was reconstructed in (1) samples with fewer species but the same sequencing depth, (2) samples with deeper sequencing but the same number of species, (3) samples with a greater abundance of unknown species (since, trivially, the reconstructed uncharacterized genomes account for more of the abundance), and (4) samples with more skewed abundances (since the highly abundant genomes that could be successfully reconstructed accounted for most of the abundance) (**Fig. 2E**, **Supplemental Fig. 8**). Assembly-based methods were mostly unaffected by low mutation rates, but a mutation rate of 0.05 (5% of bases randomly substituted for a different base) slightly decreased the abundance of correctly reconstructed and assigned genomes (mean 7% decrease in reconstructed abundance from a 2% to 5% substitution rate). This is consistent with 5% representing the intra-species average nucleotide identity threshold(47). Thus, while several parameters can impact assembly-based methods, these methods were able to recover novel genomes, particularly when the genome coverage was at least 11x to 23x (the optimal threshold for separating non-recovered from recovered genomes across methods and datasets).

In recent years, many taxonomic profilers have added the option to predict the proportion of unknown abundance in a sample, helping researchers estimate the amount of microbial “dark matter” in the metagenomes (**Methods**). To test these estimations, we compared these estimated abundances to the true abundances of uncharacterized species. The *k*-mer methods, MetaPhlAn 3 and 4, and mOTUs3 all produced generally accurate estimates in most environments (0.36% to 0.74% increase in the estimated unknown abundance for every 1% increase in the true unknown abundance) (**Fig. 2F**, **Supplemental Fig. 9**). Metaxa2 and the assembly-based methods vastly overestimated the abundance of uncharacterized species, usually because they were only able to place reads or MAGs at higher level taxonomies, not at precise species. MetaPhlAn 2 vastly underestimated the abundance of uncharacterized species because it does not explicitly estimate the unknown abundance; it simply records when the sum of abundances from lower level taxonomic markers does not align with the abundance of higher-level taxonomic markers.

### The bad: taxonomic profiling methods assign substantially different profiles to the same sample

Given the performance of the taxonomic assignment methods on our synthetic communities, we next assessed how the profiles assigned by different taxonomic assignment methods compared using real data. First, we found that the proportion of variance in taxonomic assignments explained by the method type was inversely associated with how characterized an environment was (PERMANOVA on genus-level BC dissimilarity; well-characterized R^2^ 0.09; moderately-characterized R^2^ 0.06-0.23; poorly-characterized average R^2^ 0.14-0.36; all p-values 0.0001; **Fig. 3A**). Thus, when there was more microbial dark matter, the taxonomic profilers agreed less. To limit the influence of different taxonomic naming by different methods, taxonomic names were converted into standardized NCBI IDs and only abundances from assignments with NCBI IDs were compared (**Methods**). Thus, the observed differences in real data should come from the methods’ different algorithms and databases, not the names of the taxa in those databases. When comparing taxonomic profiles assigned to the same sample by pairs of methods, well characterized environments showed more agreement among assigned genera than less characterized environments (well-characterized mean intersection over union (IOU, equivalently the Jaccard index) 0.32 and BC dissimilarity 0.37; moderately-characterized IOU 0.27, BC 0.54; poorly-characterized IOU 0.09, BC 0.84; **Fig. 3B-C**). *K*-mer methods (Centrifuge and Kraken 2/Bracken 2) showed the most agreement in assigned taxa due to their similar algorithms and databases. Other method types showed low agreement within and between types, and these trends held for both unweighted (intersection-over-union) and weighted (Bray-Curtis dissimilarity) metrics. When aggregated over non-human datasets, the profiles assigned by pairs of methods to the same sample were more similar at higher taxonomic levels (phylum level: IOU average over datasets 0.39, BC dissimilarity average over datasets 0.46; genus level: 0.15, 0.74; species level: 0.08, 0.86; **Supplemental Fig. 10**). At the species level, the three groups of methods showed more agreement with each other than with methods from other groups (**Supplemental Fig. 10**). This suggests that, to some extent, the algorithm impacts the type of classifications made by profiling methods.

**Figure 3:**
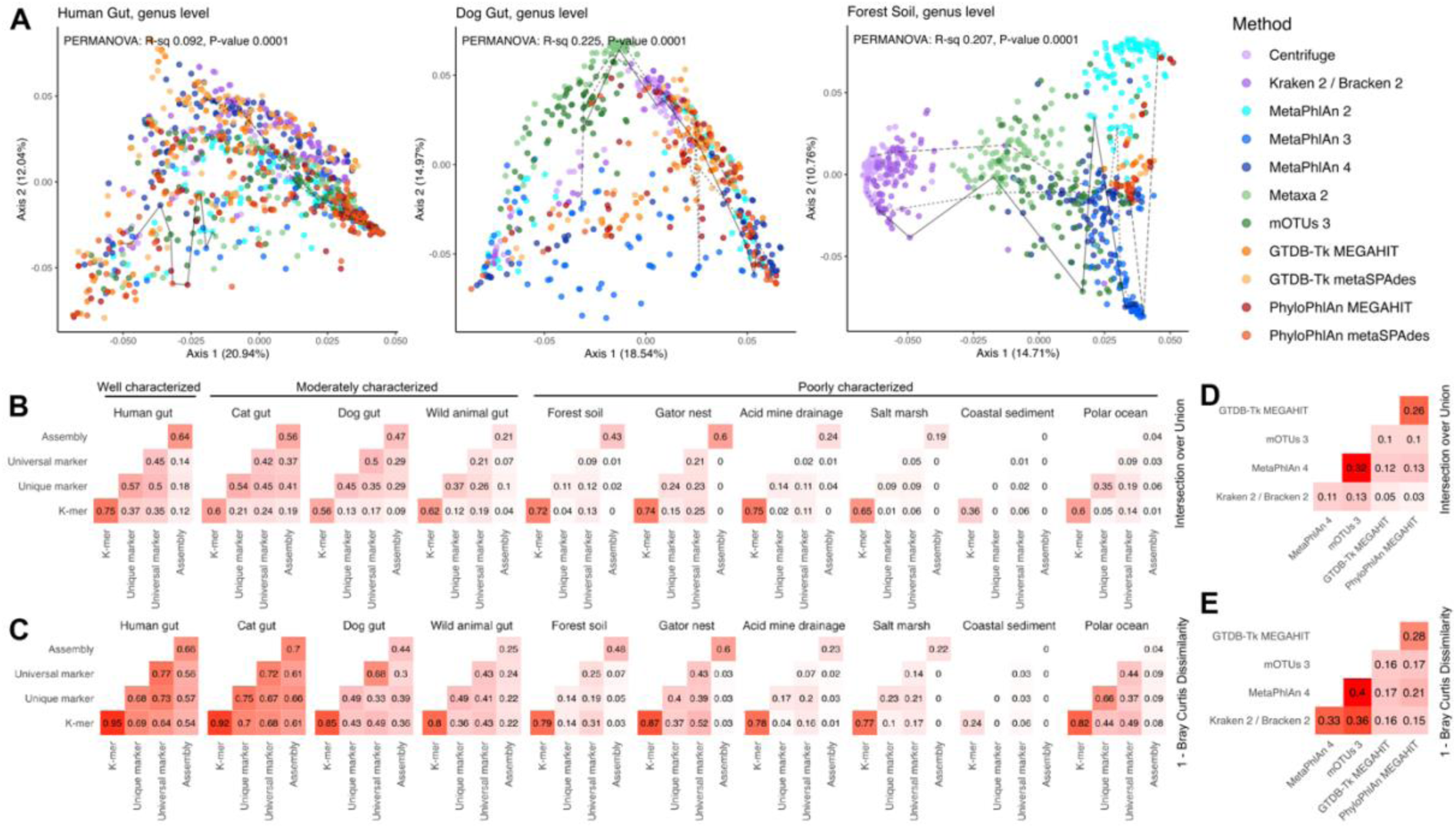
Methods assign different profiles in real environmental data. **A.** Assigned taxonomic profiles depended heavily on the method in uncharacterized environments. PCoA of Bray-Curtis dissimilarity by dataset. Lines show assignments by different methods on 3 randomly selected samples per dataset. PERMANOVA R^2^ and p-values are for the dissimilarity matrix explained by method type (*k*-mer: Kraken/Bracken 2 and Centrifuge. Unique marker: MetaPhlAn 2, 3, and 4. Universal marker: mOTUs3 and Metaxa2. Assembly: MEGAHIT or metaSPAdes with GTDB-Tk 2 or PhyloPhlAn 3) **B.** Assigned profiles differed by method more in uncharacterized environments. Average intersection over union for genus assignments on the same sample by dataset. For this figure, only the NCBI-available portion of the profile was compared. The coastal sediment assembly 0s are because no MAGs were assembled. **C.** Average Bray-Curtis dissimilarity for genus assignments on the same sample by dataset. **D.** Average intersection over union for genus assignments on the same sample averaged over the non-human datasets. **E.** Average Bray-Curtis dissimilarity for genus assignments on the same sample averaged over the non-human datasets.

### The ugly: different methods produce different microbial community structures

Because the methods significantly disagreed on which taxa were present in what abundances, we next evaluated whether the methods produced the same community structures. To assess the impact of reference-based taxonomy, we also compared the methods against Simka, a reference-free method that estimates species diversity using *k*-mer composition. Each method produced moderately similar (and significantly non-random) species dissimilarity structures with each other and with the reference-independent method Simka (Mantel r 0.36-0.94 for pairs of methods averaged across all datasets; **Fig. 4a** and **Supplemental Fig. 11**). *K*-mer methods showed more agreement (species Mantel r averaged across datasets 0.94) than other pairs of reference- and assembly-based methods (Mantel r 0.36-0.77). The correlation of the inverse Simpson alpha diversities assigned by a pair of methods to samples in a dataset was weaker, particularly at the species level (Spearman correlations averaged across datasets -0.06 to 0.87; **Fig. 4b** and **Supplemental Fig. 11**). *K*-mer methods showed more agreement with each other (0.87 at the species level), and assembly methods showed slightly more agreement among themselves (0.53-0.80). Otherwise, methods showed generally weak correlations at all taxonomic levels (lower magnitudes and fewer significantly non-zero correlations than for beta diversity). Using the alternative Shannon index or the correlation in how many taxa were assigned to a sample produced similarly inconsistent results (Shannon Spearman correlation -0.08 to 0.88; species count correlation -0.15 to 0.86).

**Figure 4:**
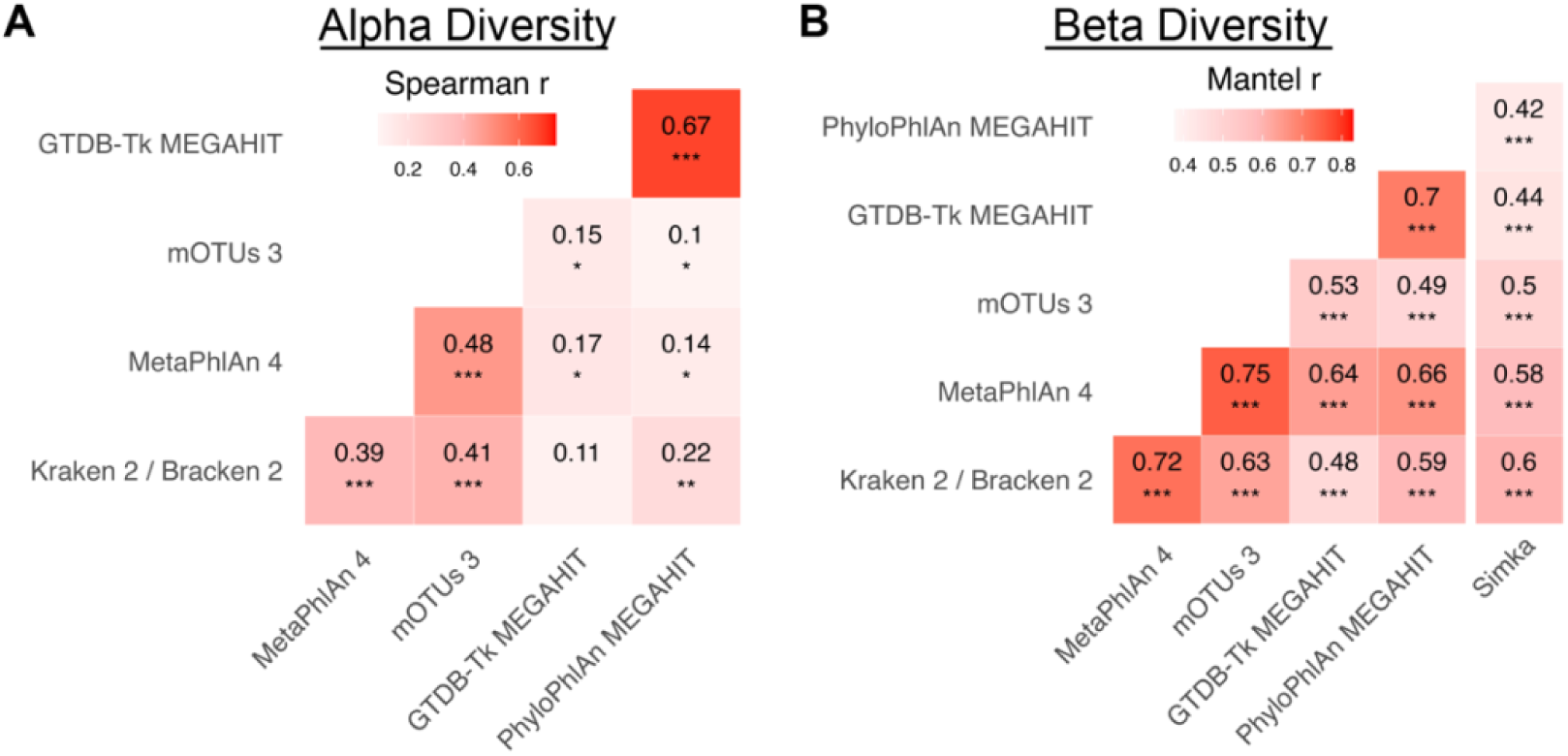
Inferred community structure varies by method. **A.** Methods agreed poorly on sample alpha diversity (inverse Simpson) aggregated over non-human datasets. An alpha diversity was computed for each sample from each method’s profile, and Spearman correlations were computed for each pair of methods for each dataset. Spearman r values are the average of Spearman r values for each non-human dataset. Significance is from a one-sided t-test against the null hypothesis that the true average Spearman r for the datasets is at most 0. **B.** Methods moderately agreed on community similarity structures (beta diversity) over non-human datasets. A Bray-Curtis dissimilarity matrix for samples in each dataset was calculated from each method, and pairs of matrices were compared. Mantel r values are the average of the Mantel r values for each non-human dataset. Significance is from a one-sided t-test against the null hypothesis that the true average Mantel r for the datasets is at most 0. Stars represent significance (p-value < 0.05, <0.01, <0.001). For both panels, all taxa assigned to a sample (NCBI-available or not) were used to compute the Bray-Curtis dissimilarity matrix or inverse Simpson metric.

The predicted abundance of unknown taxa in real community samples also differed significantly by method with very high predicted unknown abundances at low taxonomic levels (**Supplemental Fig. 2**). In particular, at the species level, all methods estimated that more than 50% of samples had more than 80% uncharacterized species abundance. As in the simulated data, MetaPhlAn 2 predicted lower unknown abundances than other methods, and the assembly methods and Metaxa2 predicted high unknown abundances at low taxonomic levels. For all methods, the proportion of the abundance considered unknown increased at lower taxonomic levels with MetaPhlAn 2, Metaxa2, and mOTUs3 displaying the largest increases in unknown abundance between the phylum and species levels.

### The okay: downstream analysis shows moderate agreement on the most significant effects

It is possible that different taxonomic profilers could identify the same environmental properties as important to microbial abundances even if the profilers disagreed on what taxa were present. To assess this possibility, we first examined whether the different profilers found similar amounts of community composition change associated with environmental variables using a univariate PERMANOVA test for each available metadata variable. For the two well- or moderately-characterized environments with available metadata, the taxonomic assignment methods produced similar PERMANOVA estimates (average range of R^2^ across methods per variable 0.07) and p-values (**Fig. 5A** and **Supplemental Fig. 12A**). By contrast, the less-characterized environments showed significant variability in their PERMANOVA results (average range of R^2^ across methods per variable 0.30). This was partially explainable by the fact that the acid mine drainage and salt marsh datasets only had 11 and 20 samples respectively. However, even excluding these datasets, the R^2^ range across methods still averaged 0.21 in uncharacterized environments. Since this range was often larger than the difference in R^2^ between variables, a variable that explains the most taxonomic variance with one method will not necessarily be the variable that will explain the most variance with another method.

**Figure 5:**
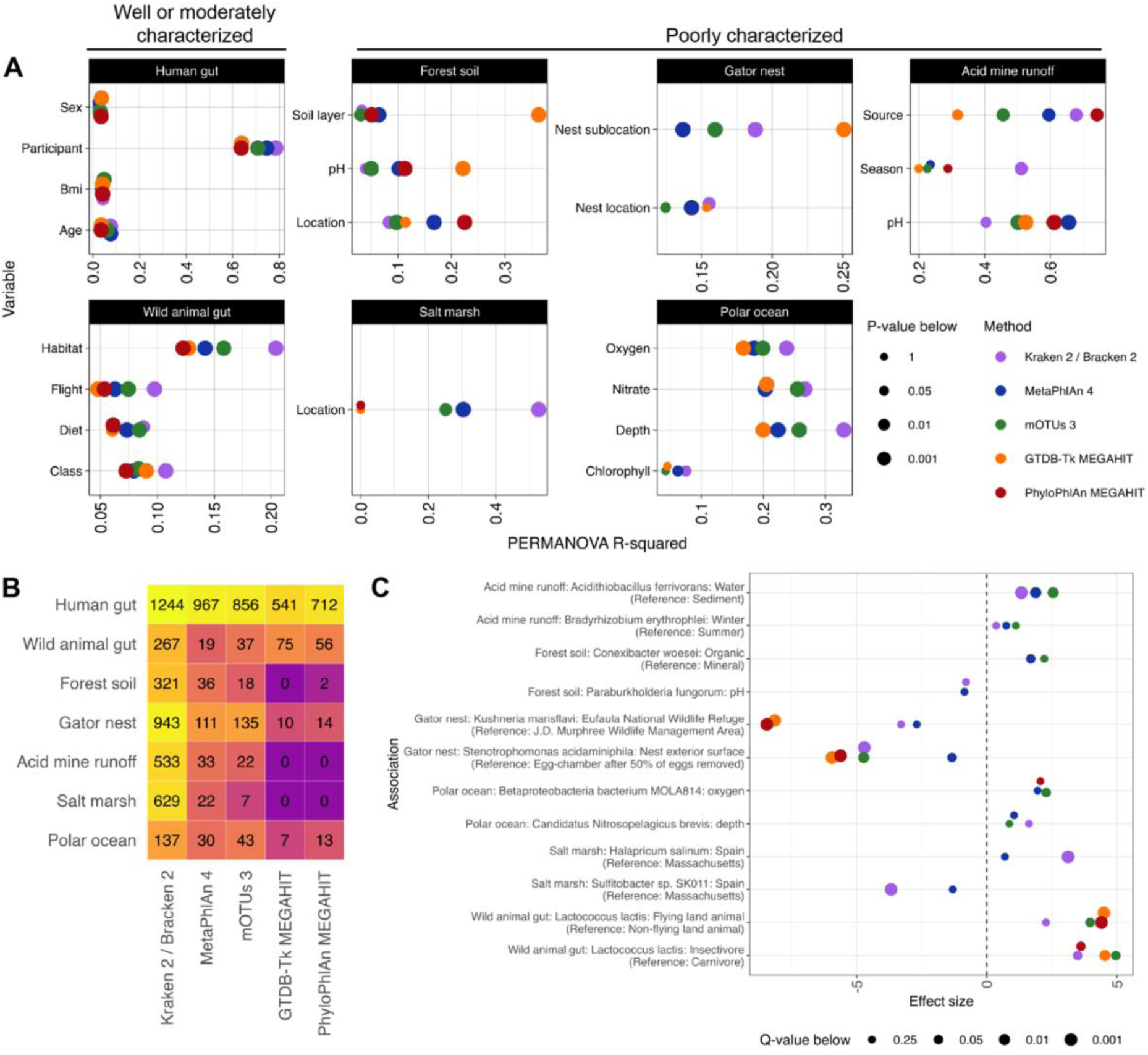
Methods show improved agreement in identifying important microbial associations. **A.** PERMANOVA consistency differs by dataset. A PERMANOVA of Bray-Curtis dissimilarity explained by sample metadata was performed for each dataset for each method. Human associations were controlled for the subject; all other associations were univariate. **B.** The number of significant (q-value < 0.25) species-metadata associations as determined by MaAsLin 2 differs by dataset and by method. **C.** The most commonly significant differential abundance results show similar effect sizes. For each non-human dataset, the log2 fold changes for the two species-metadata associations identified as significant by the most methods. A MaAsLin 2 effect size is only shown for a method if the q-value from its profile’s association was less than 0.25.

We also used MaAsLin 2(48) to perform differential abundance testing on each metadata-species pair. In the resulting associations, different methods find different numbers of significant associations, and methods that assign more species to a sample (e.g., Kraken 2/Bracken 2) tend to produce more significant associations despite the false discovery rate correction (**Fig. 5B** and **Supplemental Fig. 12C**). In particular, assembly-based methods were less likely to discover significant associations, an expected result given that they are only characterizing the most abundant features. Additionally, there was poor agreement on which metadata-species associations were significant among methods (0%-40% intersection over union for pairs of methods averaged over the datasets; **Supplemental Fig. 12B**). However, the metadata-species associations most commonly found to be significant generally had similar effect sizes across the methods (**Fig. 5C** and **Supplemental Fig. 12D**). Thus, while methods assign both different species and different abundances, when methods actually identify the same species and MaAsLin identifies that species as significant, the association is often similar across methods.

## Discussion

Environmental microbial communities play many important roles in ecological and human-relevant settings, but they present challenges for taxonomic characterization due to high levels of diversity and the relative lack of previous characterization. Here, by evaluating state-of-the-art taxonomic classification methods on a variety of real and simulated metagenomes, we show that taxonomic classification performance is often quite context-dependent. Overall, our results suggest that the choice of taxonomic assignment method should depend on the research questions, and, given the current state of the field, employing multiple methods (reference- and assembly-based) is likely the best approach.

Over a wide range of sample parameters, MetaPhlAn 4 and mOTUs3 recovered the most accurate taxonomic profiles for the portion of species that could be evaluated (species with taxonomy recorded in NCBI; **Fig. 2B-C**; **Methods**). Relative to other reference-based taxonomic classifiers, MetaPhlAn 4 and mOTUs3 both incorporate substantial numbers of MAGs (many of these from environmental sources) in their databases. While species represented by these MAGs account for a large proportion of environmental communities, non-isolate genomes are rarely included in curated databases, so supplementing these databases with other genome sources represents a significant step forward for environmental research. The incorporation of MAGs into species bins with existing species labels appeared to improve recall substantially (**Supplemental Fig. 4-6**). Likely, this was because the MAGs improved marker gene identification, allowing the methods to better generalize beyond isolate reference genomes when identifying the high quality SGBs swapped for RefSeq isolate genomes in the evaluation (**Methods**). Thus, when analyzing environmental samples (especially samples with low sequencing depth or very high diversity), MetaPhlAn 4 and mOTUs3 appear to be ideal choices for profiling the “known” component of the community. As their databases of environmentally sourced genomes expand (in part due to work on environmental microbiomes by groups like the National Microbiome Data Collaborative), these methods will likely continue to improve.

By contrast, a previous evaluation with mock communities created only from NCBI-available genomes found that Kraken2 with a small threshold performs best in environmental settings(49). However, most of the abundance in environmental samples is estimated to come from unknown species (**Supplemental Fig. 2**), so evaluating the methods on communities with significant uncharacterized abundances can better approximate a real investigation. This would explain the divergent results, since we mainly focused on uncharacterized microbes in our synthetic communities.

When evaluating assembly methods, various measures of genome coverage were, as expected, the most important factors for producing accurate profiles and recovering previously uncharacterized genomes. Deeper sequencing, fewer species, and (when using abundance-weighted metrics) higher abundance skew were all positively associated with accurate genome reconstruction. Practically, sequencing depths of less than 7.5 giga-base pairs resulted in almost no genomes reconstructed that could be placed accurately at the species level. Despite their challenges, the assembly-based methods, as expected, allowed us to recover a portion of the “unknown” fraction of the community (**Fig. 2E**). Although not tested for in this analysis, assembled genomes would presumably allow for further interrogation of the functional capacity of existing and novel species, as previously reported in environmental and human studies(18, 50).

On real environmental datasets, accuracy could not be directly assessed, but the profiles produced by different methods often clustered more by the profiling method and profiling algorithm than the underlying sample itself. Additionally, the lack of consistency between profilers was more extreme in less characterized environments, suggesting that much of the disagreement arises from different assignments on the uncharacterized species abundance. When evaluating community structures, methods showed more agreement on beta diversity than alpha diversity, consistent with slightly lower alpha diversity accuracy in simulated profiles. Likewise, all methods left the vast majority of species abundance in most samples unclassified, indicating that much work remains to fully characterize these communities. When identifying associations between a microbial community and its environmental parameters (e.g., pH or oxygen content), the profilers varied substantially in their inferences on all but the most significant associations. The datasets that produced the most inferential variability were datasets that had few samples, shallow sequencing, and uncharacterized environments (e.g., salt marsh and acid mine runoff). Still, even datasets with many samples (e.g., North American forest soils) or deep sequencing (e.g., polar oceans) showed substantial variability in PERMANOVA results. While these results are worrisome for generalizing biological findings, we were encouraged by the fact that the most significant differential abundance relationships always had the same sign and typically had similar effect sizes. Taken together, these results indicate the need for caution when analyzing such datasets and suggest that multiple lines of evidence should be employed to strengthen conclusions when much of the community consists of microbial dark matter.

## Conclusions

Here, we assess the impact of sequencing depth, degree of prior characterization, and diversity in environmental communities on a variety of state-of-the-art metagenomic analysis methods. While all methods performed similarly at high taxonomic ranks, newer marker-based methods (MetaPhlAn 4 and mOTUs3) outperformed others at lower taxonomic levels. As expected, assembly-based methods were most impacted by sample complexity and sequencing depth. Reference-based methods exhibited lower accuracy at higher taxonomic ranks when a greater proportion of uncharacterized species was present (even when high-level taxonomies for those species were available). Still, reference-based methods often recovered accurate community-level information, including within-sample diversity and between-sample similarity. In real environmental data from soil and ocean settings, taxonomic profiles assigned to the same sample showed substantial method dependence with little agreement at lower taxonomic levels, regardless of the metric used (weighted or unweighted). What similarity did exist depended strongly on the type of algorithm employed, with similar methods demonstrating greater agreement. However, as in simulated data, even disparate methods agreed substantially on between-sample similarity structures and to a lesser extent on estimates of the relationships between environmental parameters and microbial profiles in those environments.

Ultimately, this analysis suggests that a reasonable taxonomic workflow for an environmental setting would involve (1) sequencing samples as deeply as is feasible, (2) assembling and phylogenetically placing genomes in order to study previously uncharacterized microbes, (3) taxonomically profiling the samples with a reference-based method that incorporates environmental MAGs (MetaPhlAn 4 or mOTUs3) to characterize the many microbes whose genomes cannot be assembled, and (4) potentially taxonomically comparing the samples with a *k*-mer approach or a taxonomy free approach if community structure metrics are key goals. Furthermore, if computationally feasible, assembled genomes from step (2) should be added to the databases used in steps (3) and (4) to allow for further characterization of the community. Associations found between environmental parameters and the microbial community should be regarded carefully, and ensembling multiple methods may enable more robust inference. As databases incorporate more environmentally sourced genomes, the accuracy of taxonomic profilers in environmental settings will likely continue to improve, and our understanding of these communities will grow along with it.

## Methods

### Environmental dataset selection and preprocessing

#### Data selection

Environmental datasets were selected to represent a diverse set of microbial communities(23–31) (**Supplemental Table 1**). All included datasets used Illumina sequencing platforms for whole genome metagenomic sequencing, and raw reads were publicly available from the European Nucleotide Archive. All reads except the gator nest soil samples were paired-end. For the North American forest soil dataset, samples ERR1346885 and ERR1346886 were excluded since they were the only single-end samples in an otherwise paired-end dataset. In the gator nest soil dataset, sample SRR9697565 was excluded since it was the only paired-end sample in an otherwise single-end dataset. For the human gut dataset, 100 randomly selected samples from healthy controls were included. Of the salt marsh samples available from the Earth Microbiome Project, 20 were selected for having at least 500,000 reads. Of the animal stool samples available from the Earth Microbiome Project, 62 were selected for having at least 1 million reads and minimizing repeated host species. For the remaining datasets, all samples were used.

#### Metadata acquisition

Metadata for the human gut samples was obtained from the publicly available data for HMP2(31). Metadata for the North American forest soils was obtained from supplementary tables 1 and 4 of its corresponding paper. All other metadata was obtained with the ‘get_ena_file_reports.R’ script, which matched publicly available read files to their publicly available biosample information and compiled any metadata available there.

#### Data preprocessing

Raw read files were quality controlled with KneadData 0.12.0, which trims adaptors and low quality reads and performs decontamination against host genomes. The script ‘kneaddata_workflow.py’ ran ‘kneaddatà with the parameters ‘--serial --run-trf’. The cat gut samples used a contaminate database built from the felis catus genome (GCA_000181335); the dog gut samples used a contaminate database built from the canis lupus familiaris genome (GCF_014441545); and all other samples used a contaminate database built from the GRCh37 human genome.

### Simulation workflow

#### Genome curation

To create taxonomic profiles representative of the highly uncharacterized microbial communities found in environmental samples, three types of genomes were included. First, we included genomes from RefSeq (labeled “representative” or “reference”) for species listed in NCBI’s taxonomy database before the most recent method database update, GTDB-Tk’s May 11^th^ 2022 release. These genomes are hereafter referred to as “NCBI-available.” Second, we included genomes from RefSeq for species added to NCBI’s taxonomy database between the most recent method database update and when the simulations were run, May 11^th^ 2022 to December 27^th^ 2022. These genomes are hereafter referred to as “new NCBI.” Third, we included new SGBs without species-level taxonomic placements (hereafter “new SGB”) curated from the relevant sample types in the real data we analyzed. This strategy was chosen to balance the goals of (1) having some well-characterized species that could be identified fully, (2) curating enough uncharacterized genomes that most of the sample (even with hundreds of species) could consist of unknown species, and (3) ensuring that the unknown genomes were high quality and relevant to the environment of interest. For each simulated environment, a list of relevant NCBI-available genomes was curated with ‘makeProfiles.R’ by ranking the species identified in relevant datasets according to their average abundance across all taxonomic assignment methods. For the soil profiles, the relevant dataset was the North American forest soil dataset; for the ocean profiles, the relevant dataset was the Tara Polar dataset; and for the animal gut profiles, the relevant datasets were the cat gut, dog gut, and wild animal gut datasets. SGBs were curated by clustering MEGAHIT-generated MAGs into SGBs as previously described(18) (**Assembly-based methods**). The representative genome for each SGB was then assigned a taxonomy with GTDB-Tk (**Assembly-based methods**). Because GTDB-Tk’s taxonomy differs from NCBI’s taxonomy, for each SGB’s assignment, the lowest level assigned taxon recognized by NCBI was determined by an Entrez lookup to the NCBI taxonomy database(51). First, if the SGB’s assignment did not include an NCBI-recognized species, it was marked as an unknown species of the lowest taxonomic group for which it had an NCBI-recognized taxa. Up to 10 new SGBs from each genus were allowed in each profile chosen based on a ranking of the completeness minus five times the contamination. Otherwise, if (1) the new SGB had an NCBI-recognized species assignment, (2) that species was already present in the list of NCBI-available genomes, and (3) the SGB was more than 90% complete with less than 5% contamination, the RefSeq genome was replaced with the SGB under the assumption that the SGB would better exemplify a strain of the species found in the environment. We used the GTDB-Tk 2 assignments of these SGBs as the true assignments rather than the PhyloPhlAn 3 assignments due to GTDB-Tk 2’s high reported accuracy(44, 52) and the fact that this would not bias the evaluation towards the MetaPhlAn suite of tools that rely on a similar database to PhyloPhlAn 3. This procedure of using the GTDB-Tk assignments for SGBs as the true taxonomic assignment risks biasing the results in favor of GTDB-Tk, but, because assignments were usually only retained at non-species taxonomic levels, this did not appear to happen except possibly in upper-level taxonomic evaluation of samples with 100% unknown proportions (**Fig. 2C**). Full profiles are available at https://github.com/WillNickols/environmental_methods_comparison.

#### Parameter selection

We evaluated how the accuracy of the taxonomic assignment methods varied with potentially relevant sample parameters: the number of species included in a sample (species count), the sequencing depth (sample size), the proportion of new NCBI genomes and new SGBs (unknown species), the random point mutation rate for all input genomes (mutation rate), the variance in genome size (genome size skew), and the variance of the log normal distribution underlying taxonomic abundances (abundance skew). In each simulated profile, one parameter was varied while the others were held constant at a default value. To ensure we had enough uncharacterized genomes for mostly-unknown samples with many species while still capturing the diversity of environmental samples, the number of species per sample was 75, 150, 300, or 600 with a default of 300. Consistent with the range of sequencing depths observed in real data (**Fig. 1e**), the sequencing depths were 0.05, 0.5, 1.5, 7.5, and 30 giga-base pairs with a default of 7.5. Consistent with estimates of uncharacterized abundances in real samples (**Supplemental Fig. 2**), the proportions of unknown species (both by number and abundance) were 0, 0.25, 0.5, 0.75, and 1 with a default of 0.75 for soil and ocean samples and a default of 0.5 for animal gut profiles. The rates of random point mutations throughout genomes were 0, 0.01, 0.02, and 0.05 (the recommended species threshold(53)) with a default of 0 since these mutations do not necessarily represent accurate mutation patterns (e.g. conserved regions were not taken into account). Due to concerns about genome sizes affecting profilers differentially(46), various genome skews (**Supplemental Fig. 13**) were created by greedily matching genome sizes to a sequence of the form *(k-1)^j^/k^j^* normalized to the median of the genomes sizes. The values of *k* considered were 40 (widest spread), 100 (moderate spread), and 250 (narrow spread) with a default of 100. The widest spread and narrowest spread were always distinct from each other, but the widest and moderate spread were constrained by the availability of genomes, particularly in the animal gut simulation. Abundances were drawn from a log normal distribution and converted to relative abundances. The underlying normal distribution had parameters µ=-3 and σ equal to 0.5, 1, 2 or 4 with a default of 1.

#### Read generation

Broadly, once a list of NCBI-available, new NCBI, and new SGB genomes was curated and a set of parameters was specified, profiles were generated as inputs for CAMISIM(22), and CAMISIM was used to generate read files. First, the input genomes were split into known species (NCBI-available species in rank order of average abundance in real data) and unknown species (new NCBI and new SGB genomes in an arbitrary order). The known species list was truncated to only include a number of species equal to the sample’s total species number times the known proportion of the sample. This sorted list was then randomly split into two equal groups, interleaved, and assigned abundances from a sorted list of log normal draws. In this way, species that were the most common in the relevant real data were likely to be dominant in the simulated profiles, but their order and abundances could vary. These abundances were then converted to relative abundances such that their sum was the total known proportion of the sample. In this way, both the number and relative abundance of the known species matched the nominal value. For new NCBI and SGB genomes, genome lengths were calculated, and genomes were selected for inclusion by greedily matching genome sizes to the sequence determined by *k* after the known genomes were already inserted into the sequence. The number of new NCBI and SGB genomes included was set equal to the total number of species in the sample times the unknown proportion of the sample. These species were then assigned abundances from a log normal distribution with the same parameters as the known profile, but the abundances were then converted into relative abundances such that their sum was the total unknown proportion in the sample. If a non-zero mutation rate was specified for the sample, substitution point mutations were made at the specified rate in all known and unknown genomes for the sample. Finally, configuration files for CAMISIM were generated from these profiles, and CAMISIM was run in art_illumina-2.3.6 mode (150 base paired-end reads) with default parameters except for the number of giga-base pairs, which was set according to the sample parameters. Because we were not directly evaluating assembly methods and the gold standard assembly sections of CAMISIM required significant computing time, we removed the gold standard assembly sections from the CAMISIM code. Otherwise, CAMISIM was run unmodified.

### Reference-based methods

To avoid preferencing any particular method and because many methods were evaluated, no parameters were tuned for any method. The only parameters changed from their default values were flags to include more taxonomic information and output results in particular formats.

#### Centrifuge

Centrifuge(33) 1.0.4 was installed according to its installation instructions and was run from cleaned reads with the script ‘centrifuge_workflow.py’. This script ran ‘centrifugè in paired or unpaired mode as necessary with the default parameters. All bacterial, archaeal, and viral genomes were downloaded from Refseq on January 5, 2023, but the Centrifuge index was built with only genomes available before and including March 25, 2022 for consistency with the Kraken 2 database.

#### Kraken 2/Bracken 2

Kraken 2.1.2(34) was installed according to its installation instructions, and its standard database was built on March 25, 2022 with ‘kraken2-build --standard’. Bracken 2.6.2(35) was installed according to its installation instructions, and its default database was built on March 25, 2022 with ‘bracken-build’. Kraken 2 and Bracken 2 were run with the script ‘kraken_workflow.py’ on cleaned reads. This script ran ‘kraken2’ and ‘bracken’ in paired or unpaired mode as necessary with default parameters.

#### MetaPhlAn

MetaPhlAn 2.6.0(37) was installed according to its installation instructions with the database mpa_v20_m200. MetaPhlAn 3.0.14(38) was installed according to its installation instructions with the database mpa_v30_CHOCOPhlAn_201901. MetaPhlAn 4.beta.2(20) was installed according to its installation instructions with the database mpa_vJan21_CHOCOPhlAnSGB_202103. More recent MetaPhlAn 4 databases incorporate the MAGs generated in this study and therefore were not used. MetaPhlAn 2, MetaPhlAn 3, and MetaPhlAn 4 were run with the scripts ‘mpa2_workflow.py’, ‘mpa3_workflow.py’, and ‘mpa4_workflow.py’ respectively on cleaned reads. The ‘mpa2_workflow.py’ script ran ‘metaphlan2’ in paired or unpaired mode as necessary with default parameters; the ‘mpa3_workflow.py’ script ran ‘metaphlan’ in paired or unpaired mode as necessary with the parameters ‘--add_viruses --unknown_estimation’; and the ‘mpa4_workflow.py’ script ran ‘metaphlan’ in paired or unpaired mode as necessary with the parameters ‘--add_viruses -- unclassified_estimation --sgb’.

#### Metaxa2

Metaxa 2.2.3(36) was installed according to its installation instructions and run from cleaned reads with the default 2.2.3 database and the script ‘metaxa2_workflow.py’. This script ran ‘metaxa2’, ‘metaxa2_ttt’, and ‘metaxa2_dc’ in paired or unpaired mode as necessary with default parameters on all taxonomic levels.

#### mOTUs3

mOTUs3(19) 3.0.1 was installed according to its installation instructions and run from cleaned reads with the database db_mOTU_v3.0.1 and the script ‘mOTUs3_workflow.py’. This script ran ‘motus profilè with the flag ‘-À to report all taxonomic levels.

### Assembly-based methods

Consistent with previous approaches(18), the assembly methods consisted of contig assembly, MAG curation from contig binning, quality control, and MAG phylogenetic placement. First, either MEGAHIT 1.2.9(39) or metaSPAdes 3.15.4(40) was used to create contigs with the command ‘megahit’ (‘--min-contig-len 1500’, paired or unpaired mode as necessary) or ‘spades.py --metà (default parameters, paired mode only). Samples with over approximately 25 giga-base pairs depending on the environment were infeasible to assemble with metaSPAdes, so only MEGAHIT assembly was evaluated for these. Next, a bowtie2 (2.5.0)(54) index was built with ‘bowtie2-build’ (default parameters) from the contigs, and reads were aligned with ‘bowtie2’ (parameters ‘--very-sensitive-local’ and ‘--no-unal’). Contig read depth means and variances were calculated with ‘samtools view’ (parameters ‘-bS -F 4’; Samtools 1.16.1(55)), ‘samtools sort’ (default parameters), and ‘jgi_summarize_bam_contig_depths’ (default parameters). Contigs were binned with MetaBAT 2.15(41) using ‘metabat2’ with a minimum contig length of 1500 (‘-m 1500’). MAG abundance was determined with a by-sample approach (reads from a sample were only aligned to MAGs from that same sample) or a by-dataset approach (reads from a sample were aligned to any MAGs from that sample’s dataset) based on the analysis goal. In the by-sample approach, alignments to the original contigs were indexed with ‘samtools index’ and converted into relative abundances with the ‘checkm.py coveragè (default parameters) and ‘checkm.py profilè (‘--tab_tablè) scripts from Checkm 1.2.2(56). The total number of mapped reads was recorded with ‘samtools view -c -F 260’. In the by-dataset approach, a new bowtie2 index was created as before but from all the contigs in the dataset rather than just one sample’s contigs. The reads were then aligned, indexed, and converted into relative abundances as before. The N50 of each MAG was computed with ‘assembly-stats’ from Assembly-stats(57), and the completeness and contamination of the MAGs were assessed with Checkm2 1.0.0(42) using ‘checkm2 predict’ with default parameters. Only medium or high quality MAGs (>50% completeness and <10% contamination) were retained. MAGs were then phylogenetically placed with either PhyloPhlAn 3.0.67(43) (SGB.Jul20 database) or GTDB-Tk 2.1.0(44) (R207_v2 database) using the commands ‘phylophlan_metagenomic’ (‘-n 1 --add_ggb --add_fgb’) or ‘gtdbtk classify_wf’ (default parameters) respectively. Because PhyloPhlAn gives the closest species genome bin (SGB), genus genome bin (GGB), and family genome bin (FGB) to the MAG regardless of the actual Mash distance, the MAG was assigned to the SGB, GGB, or FGB if the Mash distance was less than 0.05, 0.15, or 0.3 respectively using previously described thresholds(18). MAGs with Mash distances of more than 0.3 to their closest FGB were left unplaced. Any phylogenetically unplaced MAGs were then grouped into new SGBs as previously described(18). This workflow was run with the scripts ‘assembly_workflow.py’ and ‘gtdbtk_workflow.py’.

### Reference independent methods

#### Simka

Simka 1.5.1(32) was installed according to its installation instructions, and the command ‘simkà was run on each dataset (‘simka_workflow.py’) for reference-independent (**Fig. 4a**) beta diversity or on all datasets together (‘simka_workflow_all.py’) to generate distances used in tSNE (**Fig. 1F**).

### Statistical analysis

#### Raw database acquisition and naming harmonization

Because different taxonomic assignment methods often used different, synonymous names for the same taxa, a conversion to a standardized naming system was necessary for F1, intersection over union, and Bray-Curtis metrics. First, a list of all taxa in each method’s database was obtained from a file created during installation (‘seqid2taxid.map’ for Centrifuge and Kraken 2/Bracken 2), a file in the source code (‘gtdb_taxonomy.tsv’ for GTDB-Tk, ‘blast.taxonomy.txt’ for Metaxa2, and ‘SGB.Jul20.txt’ for PhyloPhlAn), a conversion of a file in the source code (‘mpa_v20_m200.pkl’ for MetaPhlAn 2, ‘mpa_v30_CHOCOPhlAn_201901.pkl’ for MetaPhlAn 3, and ‘mpa_vJan21_CHOCOPhlAnSGB_202103.pkl’ fro MetaPhlAn 4), or publicly accessible files online (‘db_mOTU_taxonomy_CAMI.tsv’, ‘db_mOTU_taxonomy_meta-mOTUs.tsv’, and ‘db_mOTU_taxonomy_ref-mOTUs.tsv’ for mOTUs3). Then, all taxa from each of these lists were searched in the ‘names.dmp’ file of NCBI’s taxonomy FTP, and a numerical NCBI ID was recorded in a mapping file for each taxa if a match was found. Because this ‘names.dmp’ file maps any recognized taxonomic synonyms to the same NCBI ID, this procedure standardizes the naming as much as is feasible. For each method’s taxonomic assignment outputs, the method’s mapping file was used to convert the raw output taxonomies into standardized NCBI IDs, allowing comparison between the profiles. This naming harmonization converts the vast majority of assigned taxa into NCBI IDs in all evaluations. Furthermore, for any evaluation involving specific taxa names (F1, BC dissimilarity, IOU, PCoA, or MaAsLin), only the taxa converted to NCBI IDs were used, and their abundances were renormalized to account for the removed non-mappable taxa.

#### Evaluation of uncharacterized species

When assigning taxonomy to a genome that does not belong to a species available in NCBI, there are multiple ways a method could perform correctly or incorrectly. First, at the species level, the genome could either be correctly marked unknown or correctly assigned to a species named the method’s database but not present in the NCBI database. The assigned non-NCBI species might still be incorrect, but it is possible that the database construction involved a genome from the same species even if this species is not yet recognized by NCBI, so we cannot conclude it is incorrect. However, if the genome is assigned to any species in NCBI, that assignment would be wrong by definition. Next, even if the genome is unknown at the species level, the genome will have some upper level taxonomy present in NCBI. For concreteness, suppose the genome belongs to an NCBI-available family but to a genus and species of that family not yet in NCBI. Then, by the same reasoning as at the species level, it could be correct to assign the genus as unknown or to a non-NCBI genus in the method’s database, but it would be incorrect to assign it to any NCBI genus. However, at the family level and higher, the genome should be assigned to the correct NCBI taxon. Assigning it to any other taxon would be incorrect, and labeling it as unknown would also be incorrect, because it is known at the family level and above.

Based on these principles, we evaluated the taxonomic assignments on simulated data by, at each taxonomic level, removing the NCBI-unavailable portion of the true profile and the NCBI-unmappable portion of the assigned profile, renormalizing each of these so their abundances summed to 1, and comparing them. In this way, methods were not penalized for marking NCBI-unavailable abundance as unknown or as NCBI-unmappable taxa, but they were penalized for assigning NCBI-unavailable abundance to NCBI-mappable taxa and for not assigning NCBI-available abundance to the correct NCBI-mappable taxa. Restricting samples to their NCBI-available taxa had a negligible impact on the methods’ accuracies (**Supplemental Fig. 14**) but significantly improved consistency between methods in real data.

#### Thresholding, normalization, and uncharacterized abundance

To limit false positives from methods such as Centrifuge and Kraken 2/Bracken 2, after testing the effect of various thresholds on F1 scores, an abundance threshold of 0.05% was applied to all assigned profiles. Taxa in a sample with abundance less than 0.05% were removed, and the remaining abundance was renormalized to sum to 1. In addition to abundance directly labeled as unknown, abundance assigned to taxa with names that were blank or included “noname” or “incertae” was labeled as unknown. For assembly-based methods, the unknown abundance was computed as the sum of (1) the proportion of reads not aligning to any MAGs, (2) the abundance of low quality MAGs, and (3) the abundance of MAGs not placed at the taxonomic level of interest. Except for in the unknown abundance evaluation, all abundance labeled as unknown was removed, and the remaining abundance was renormalized.

#### Statistical environment and packages

All statistical analysis was performed in the R language (version 4.2.2)(58). The all-dataset tSNE plot (**Fig. 1D**) was generated with the package Rtsne(59) (perplexity 30) on inter-sample distances from Simka.

In the synthetic evaluation (**Fig. 2B-C** and **Supplemental Fig. 4-6, 14**), for a particular taxonomic rank, precision was calculated as the number of taxa assigned to a sample that were actually present in the sample (after thresholding, removal of NCBI-unmappable taxa, and normalization) divided by the number of taxa assigned to the sample (after thresholding, removal of NCBI-unmappable taxa, and normalization). For a particular taxonomic rank, the recall was calculated as the number of taxa actually present in a sample that were assigned to the sample (after thresholding, removal of NCBI-unmappable taxa, and normalization) divided by the number of taxa actually present in the sample (after removal of NCBI-unavailable taxa and normalization). The F1 score was the harmonic mean of precision and recall. For a particular taxonomic rank, the BC dissimilarity was calculated by comparing the abundances of taxa assigned to a sample (after thresholding, removal of NCBI-unmappable taxa, and normalization) to the abundances of taxa actually present in the sample (after removal of NCBI-unavailable taxa and normalization). The BC dissimilarities were calculated with the ‘vegdist’ function from the R package vegan (version 2.6-4)(60).

In the synthetic evaluation (**Fig. 2D** and **Supplemental Fig. 7**), for a particular dataset at the species level, the Mantel r was computed with the ‘mantel’ function from vegan on all-sample-versus-all-sample true and assigned BC dissimilarity matrices. These BC dissimilarity matrices were generated as above except without removal of NCBI-unmappable taxa. This retains all imputed diversity since the naming is irrelevant in this case. At the species level, the inverse Simpson index of a sample was computed with the ‘diversity’ function in vegan after thresholding and normalization for assigned profiles. For a particular rank and a particular dataset, the Spearman r reported is the Spearman correlation between the assigned inverse Simpson indices for all samples in the dataset and the true inverse Simpson indices for all samples in the dataset.

In the synthetic evaluation (**Fig. 2E** and **Supplemental Fig. 8**), an assembly-based method was considered to have successfully reconstructed an uncharacterized genome if (1) it produced a MAG that was at least 50% complete and less than 10% contaminated, (2) this MAG’s Mash distance to the original uncharacterized genome was less than its distances to any other input genome, and (3) the NCBI-mappable part of its assigned taxonomy was contained within the original taxonomy assigned for the uncharacterized genome. For example, the uncharacterized genome might have been assigned an NCBI-mappable taxonomy to the genus level, and if the reconstructed MAG was assigned the same NCBI-mappable taxonomy only to the family level (and no NCBI-mappable taxonomy at the genus or species level), this would still qualify as a successfully reconstructed genome. For each sample, the abundances of the successfully reconstructed genomes from the true profile (not the profiling method’s assigned abundances) were added together and analyzed.

In the real data evaluation (**Fig. 3B-E** and **Supplemental Fig. 10**), for a particular rank, the intersection over union (IOU) was calculated as the number of taxa overlapping between assignments from two methods on the same sample (after thresholding, removal of NCBI-unmappable taxa, and normalization) divided by the total number of unique taxa assigned to the sample by the two methods (after thresholding, removal of NCBI-unmappable taxa, and normalization). For a particular taxonomic rank, the BC dissimilarity was calculated by comparing the abundances of taxa assigned to a sample by two methods (after thresholding, removal of NCBI-unmappable taxa, and normalization). The BC dissimilarities were calculated with the ‘vegdist’ function from vegan. PCoAs (**Fig. 3A**) were generated from the dataset BC dissimilarity matrices with the ‘capscalè function from vegan. PERMANOVAs were run using the ‘adonis2’ function from vegan with 9999 permutations.

In the real data evaluation (**Fig. 4** and **Supplemental Fig. 11**), for a particular dataset at the species level, the Mantel r was computed with the ‘mantel’ function from vegan on pairs of all-sample-versus-all-sample BC dissimilarity matrices as generated above (except without removal of NCBI-unmappable taxa). One Mantel r was computed for each pair of methods on each dataset, and the displayed values are the (unweighted) average across non-human datasets. A p-value was generated from a one-sided t-test against the null hypothesis that the true average Mantel r across the datasets for a pair of methods is at most 0 (n=9 datasets). At the species level, the inverse Simpson index of an assigned profile was computed with the ‘diversity’ function in vegan after thresholding and normalization. For a particular dataset at the species level, the Spearman r reported is the Spearman correlation between inverse Simpson indices for all samples in the dataset as assigned by two methods. One Spearman r was computed for each pair of methods on each dataset, and the displayed values are the (unweighted) average across non-human datasets. A p-value was generated from a one-sided t-test against the null hypothesis that the true average Spearman r across the datasets for a pair of methods is at most 0 (n=9 datasets).

In the real data evaluation (**Fig. 5A** and **Supplemental Fig. 12A**), for a particular dataset at the species level, PERMANOVAs were run using the ‘adonis2’ function from vegan with 9999 permutations to estimate the variance in Bray-Curtis dissimilarity matrices explained by sample metadata. The Bray-Curtis dissimilarity matrices were the same as those used in the Mantel testing. Human associations were controlled for the subject (except when evaluating the effect of subjects themselves); all other associations were univariate.

In the real data evaluation (**Fig. 5B** and **Supplemental Fig. 12B-D**), for a particular dataset at the species level, differential abundance testing (log2 fold change) was performed with MaAsLin 2(48) (Microbiome Multivariable Associations with Linear Models), which employs a multivariate linear model to measure β-coefficient effect sizes for feature-covariate pairs. MaAsLin 2 also reports Benjamini–Hochberg false discovery rate adjusted q-values for metadata-microbiome feature pairs. Input taxonomic profiles were thresholded, had NCBI-unmappable taxa removed, and were subsequently renormalized. Human associations were controlled for the subject (except in the pairwise subject tests); all other associations were univariate. For categorical variables, MaAsLin selected a default category and performed pairwise testing with all other categories. The large number of significant human associations is due to the many pairwise tests with the subject as the explanatory variable.

## Declarations

### Ethics approval and consent to participate

Not applicable.

### Consent for publication

Not applicable.

### Availability of data and materials

The accessions to the datasets analyzed during the current study can be found in the Supplemental Table 1. By-sample taxonomic profiles, metadata, generated intermediate files, and code are available in the “environmental_methods_comparison” repository at https://github.com/WillNickols/environmental_methods_comparison.

## Authors’ contributions

K.N.T. and C.H. conceived the project. W.A.N. and K.N.T. determined the simulation methods and reference datasets. W.A.N. preprocessed the reference datasets. W.A.N., Y.Z., E.M.H., E.A.F., and K.N.T. designed and performed the evaluation. W.A.N., L.J.M., A.W., and K.N.T. wrote the benchmarking scripts. W.A.N. and K.N.T. analyzed the data, produced the figures, and prepared the supplementary files. W.A.N. and K.N.T. wrote the manuscript. J.T.N., F.A., M.P., E.M.H., L.H.N., and C.H. proposed essential suggestions for this study and edits for the manuscript. N.S., C.H., and K.N.T. supervised the entire project. All authors read and approved the final manuscript.

## Acknowledgements

The computations in this paper were run in part on the FASRC Cannon cluster supported by the FAS Division of Science Research Computing Group at Harvard University.

