## Supplement for "Evaluating metagenomic analyses for undercharacterized environments: what’s needed to light up the microbial dark matter?"

### Diverse environmental datasets were curated for realistic comparisons

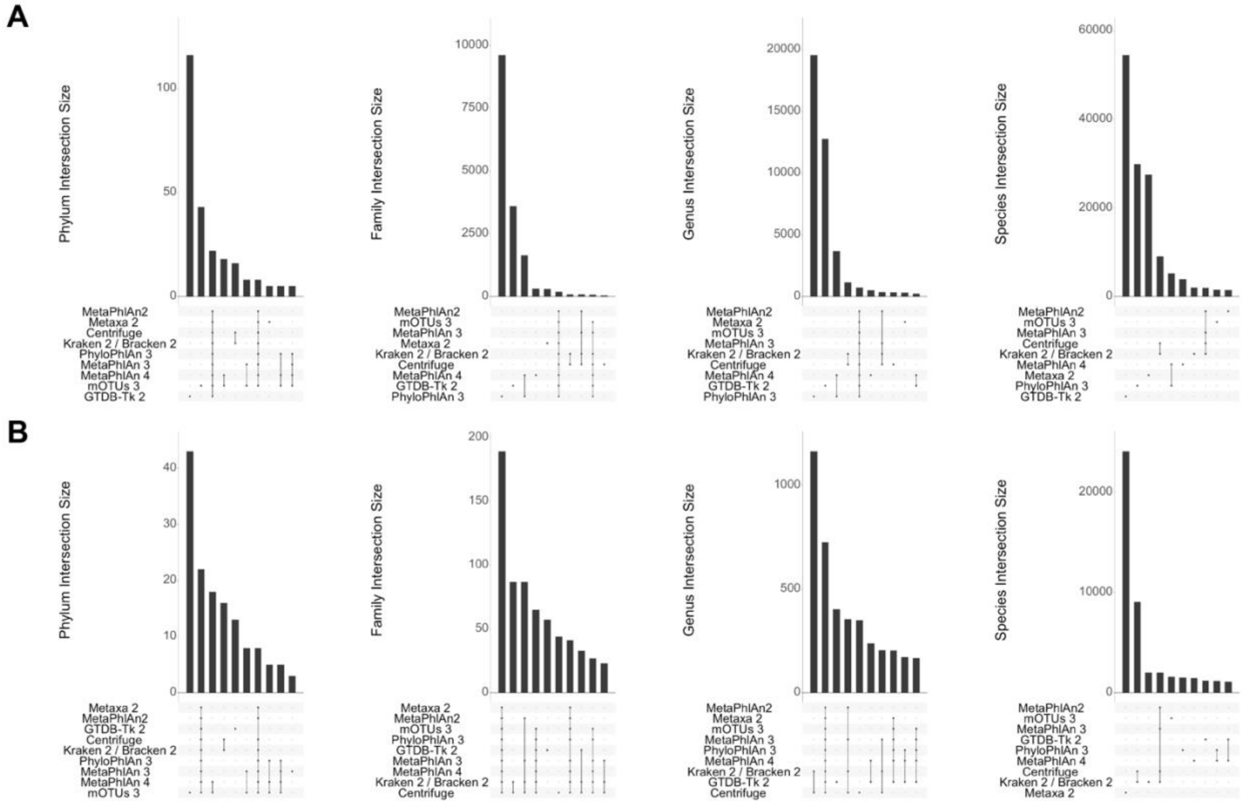

**Supplemental Figure S1: Overlap of Profiling Method Databases. A.** Number of taxonomic groups overlapping between the raw databases for each method with NCBI-harmonized naming where possible. **B.** Number of taxonomic groups overlapping between databases when restricted to NCBI-recognized taxa.

**Supplemental Table 1: Dataset Sample Overview**

| Dataset | PMID, Project ID | Location | Sample Count | Pairing | Raw reads | Mean raw reads | Cleaned reads | Mean cleaned reads |
|---|---|---|---|---|---|---|---|---|
| Acid Mine Runoff | 32785287, PRJNA540505 | Vermont, USA | 11 | Paired | 17,315,932-50,824,640 | 31,545,991 | 12,488,692-45,175,062 | 25,827,286 |
| Cat Gut | 35467389, PRJNA758898 | Auburn, AL | 16 | Paired | 54,533,852-252,204,562 | 112,935,059 | 23,611,710-244,088,713 | 94,766,490 |
| Coastal Sediment | 28793929, PRJNA322450 | Baltic Sea | 12 | Paired | 6,476,510-10,117,920 | 7,831,218 | 4,206,042-6,573,201 | 5,301,153 |
| Dog Gut | 32303546, PRJEB34360 | Leicestershire, UK | 62 | Paired | 16,558,694-125,066,282 | 86,092,539 | 14,387,659-111,409,761 | 74,846,433 |
| North American Forest Soils | 30258172, PRJEB12502 | North America | 90 | Paired | 1,057,594-390,062,486 | 78,371,093 | 401,303-358,421,172 | 68,668,864 |
| Gator Nest Soil | 32424717, PRJNA554694 | Southern USA | 34 | Single | 2,410,326-55,609,126 | 9,590,638 | 1,476,312-52,272,136 | 8,897,005 |
| HMP2 Human Controls | 31142855, PRJNA398089 | USA | 100 | Paired | 4,827,544-42,083,960 | 25,853,526 | 4,827,544-42,083,960 | 25,853,526 |
| Salt Marsh | 36443458, PRJEB42019 | MA, USA / Huelva, Spain | 20 | Paired | 593,676-150,598,352 | 13,876,036 | 277,034-70,852,992 | 6,494,136 |
| Tara Polar | 26029378, PRJEB9740 | Arctic oceans | 41 | Paired | 263,756,272-482,084,694 | 348,900,477 | 234,662,116-424,682,983 | 302,955,853 |
| Wild Animal Guts | 36443458, PRJEB42019 | USA & Malawi | 62 | Paired | 4,034,736-125,505,304 | 40,264,659 | 1,874,146-58,570,825 | 18,432,025 |

**Supplemental Table 2: Dataset Sample Overview**

| Method | Version | Database | Method Type |
|---|---|---|---|
| Centrifuge | 1.0.4 | RefSeq bacterial, archaeal, viral (March 25, 2022) | *K*-mer |
| Kraken 2/Bracken 2 | 2.1.2/2.6.2 | --standard (RefSeq bacterial, archaeal, viral, human; March 25, 2022) | *K*-mer |
| MetaPhlAn 2 | 2.6.0 | mpa_v20_m200 | Unique marker genes |
| MetaPhlAn 3 | 3.0.14 | mpa_v30_CHOCOPhlAn_201901 | Unique marker genes |
| MetaPhlAn 4 | 4.beta.2 | mpa_vJan21_CHOCOPhlAnSGB_202103 | Unique marker genes |
| Metaxa2 | 2.2.3 | 2.2.3 installed database | Universal marker genes |
| mOTUs3 | 3.0.1 | db_mOTU_v3.0.1 | Universal marker genes |
| MEGAHIT/PhyloPhlAn 3 | 1.2.9/3.0.67 | SGB.Jul20 | Assembly |
| MEGAHIT/GTDB-Tk 2 | 1.2.9/2.1.0 | R207_v2 | Assembly |
| metaSPAdes/PhyloPhlAn 3 | 3.15.4/3.0.67 | SGB.Jul20 | Assembly |
| metaSPAdes/GTDB-Tk 2 | 3.15.4/2.1.0 | R207_v2 | Assembly |

### Taxonomic profiling methods differ in accuracy on simulated environmental profiles

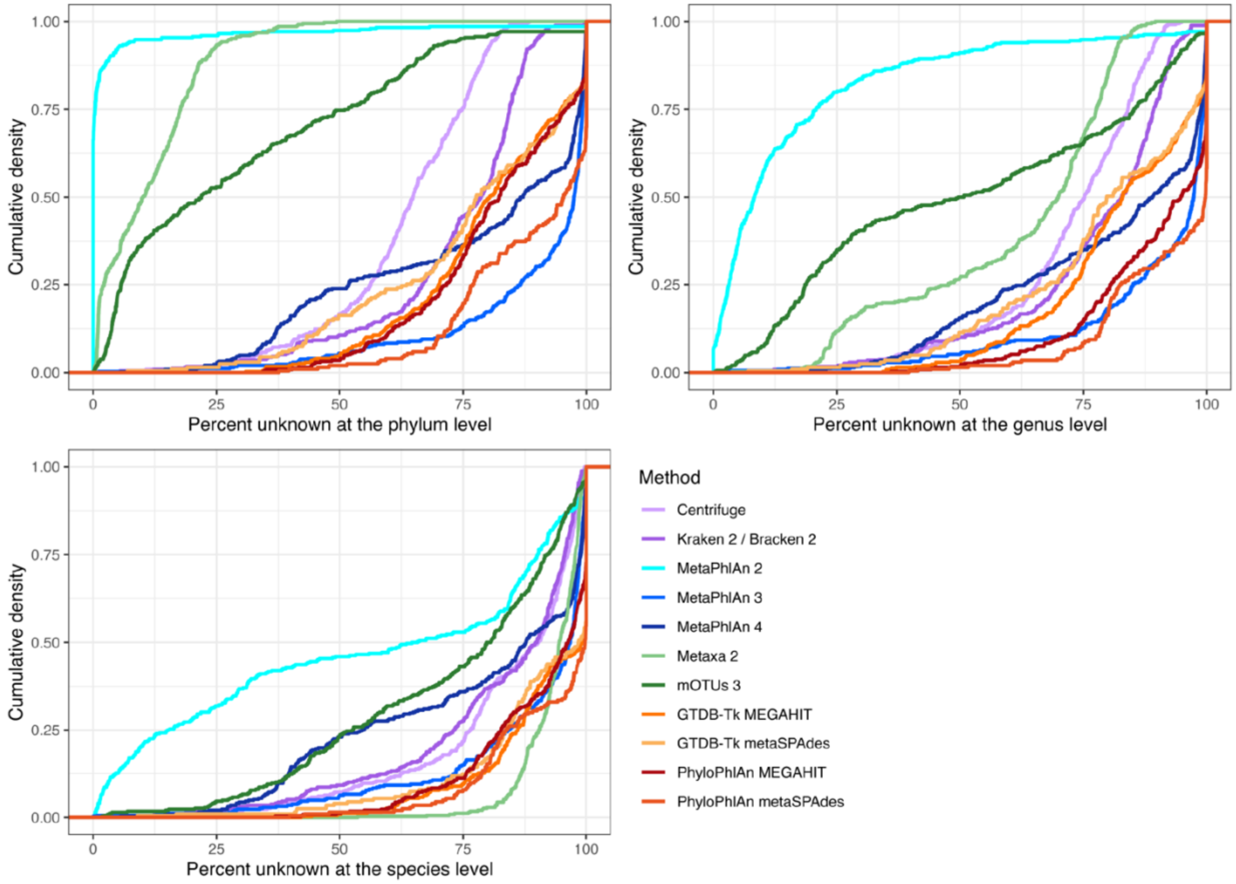

**Supplemental Figure S2: Empirical cumulative density over all real non-human samples.** For each method, for each sample, the percent of the abundance labeled unknown or left unclassified was calculated. A line that intersects a point (X, Y) on the plot indicates that the proportion of the samples with less than X% unknown was Y.

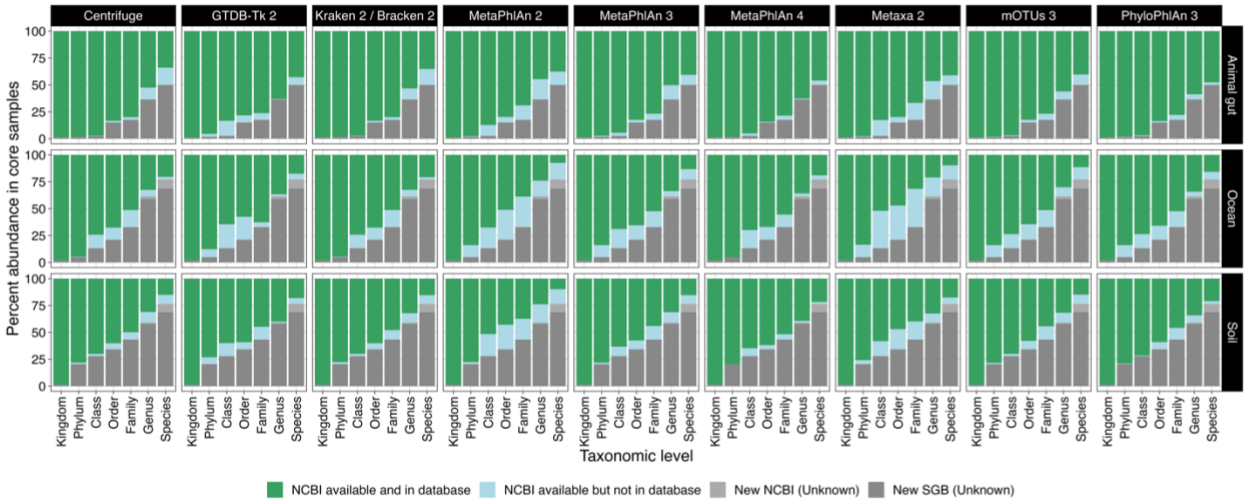

**Supplemental Figure S3: Simulation taxa present in method databases.** In core samples, the abundance of taxa with historically available NCBI identifiers in the method's database (NCBI available and in database), historically available NCBI identifiers not in the method's database (NCBI available but not in database), NCBI identifiers added since the most recent method database update (new NCBI), or no NCBI identifier (new SGB).

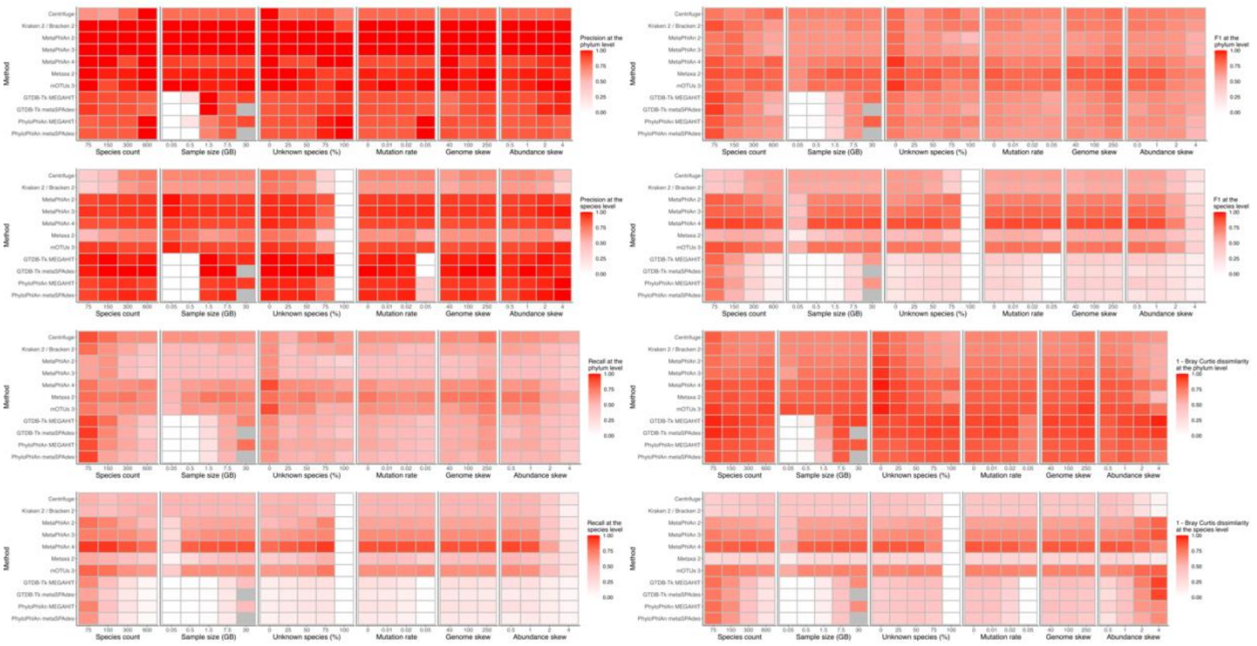

**Supplemental Figure S4: Precision, recall, F1, and Bray-Curtis dissimilarity relative to the true animal gut profile.** Metrics by sample parameter for the simulated animal gut dataset (averaged over 5 replicates per parameter setting). Only one parameter at a time was changed from the core value. Gray boxes indicate that the setting was computationally infeasible to run.

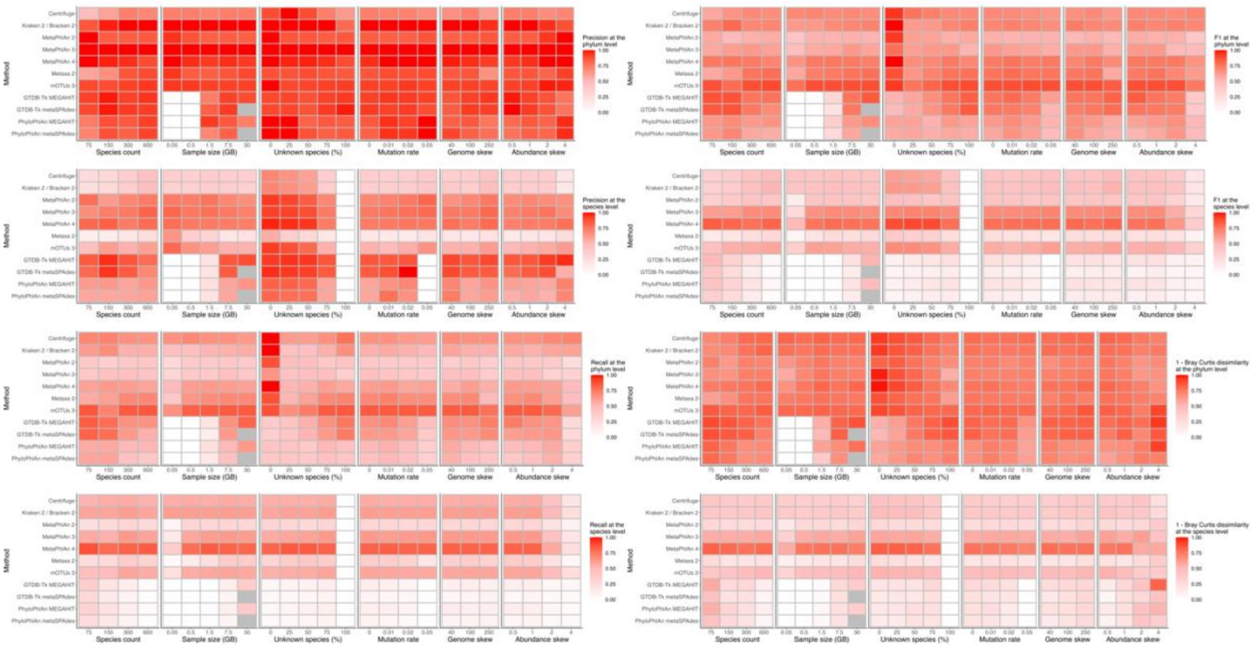

**Supplemental Figure S5: Precision, recall, F1, and Bray-Curtis dissimilarity relative to the true ocean profile.** Metrics by sample parameter for the simulated ocean dataset (averaged over 5 replicates per parameter setting). Only one parameter at a time was changed from the core value. Gray boxes indicate that the setting was computationally infeasible to run.

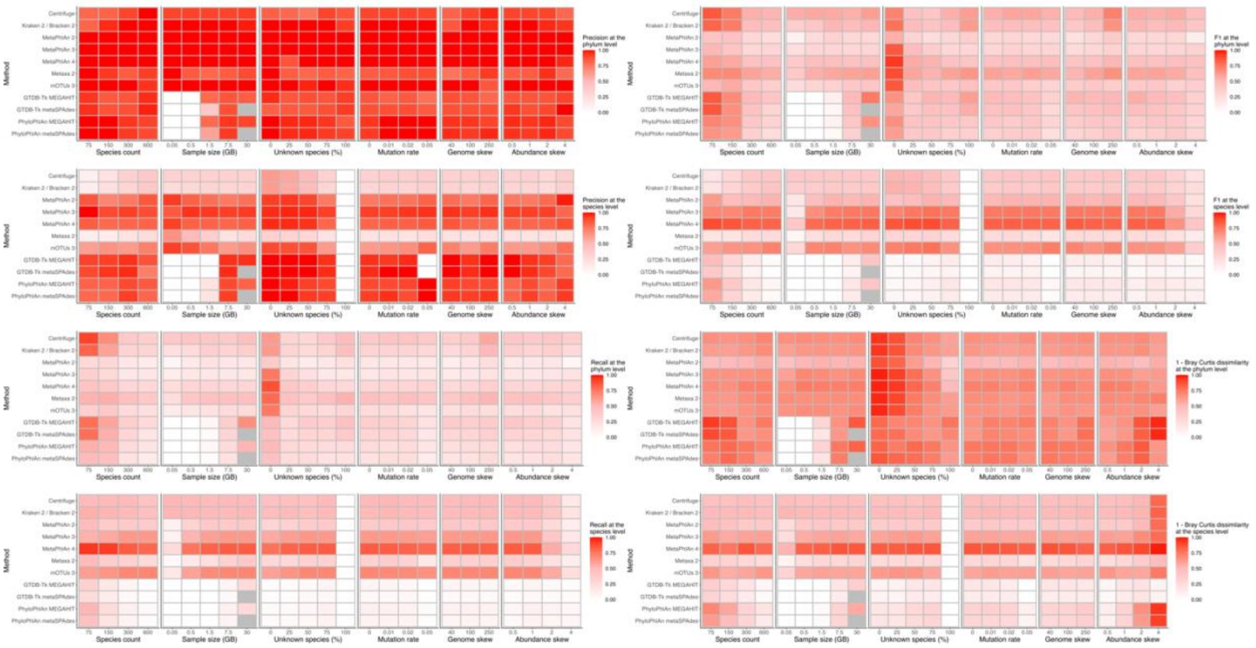

**Supplemental Figure S6: Precision, recall, F1, and Bray-Curtis dissimilarity relative to the true soil profile.** Metrics by sample parameter for the simulated soil dataset (averaged over 5 replicates per parameter setting). Only one parameter at a time was changed from the core value. Gray boxes indicate that the setting was computationally infeasible to run.

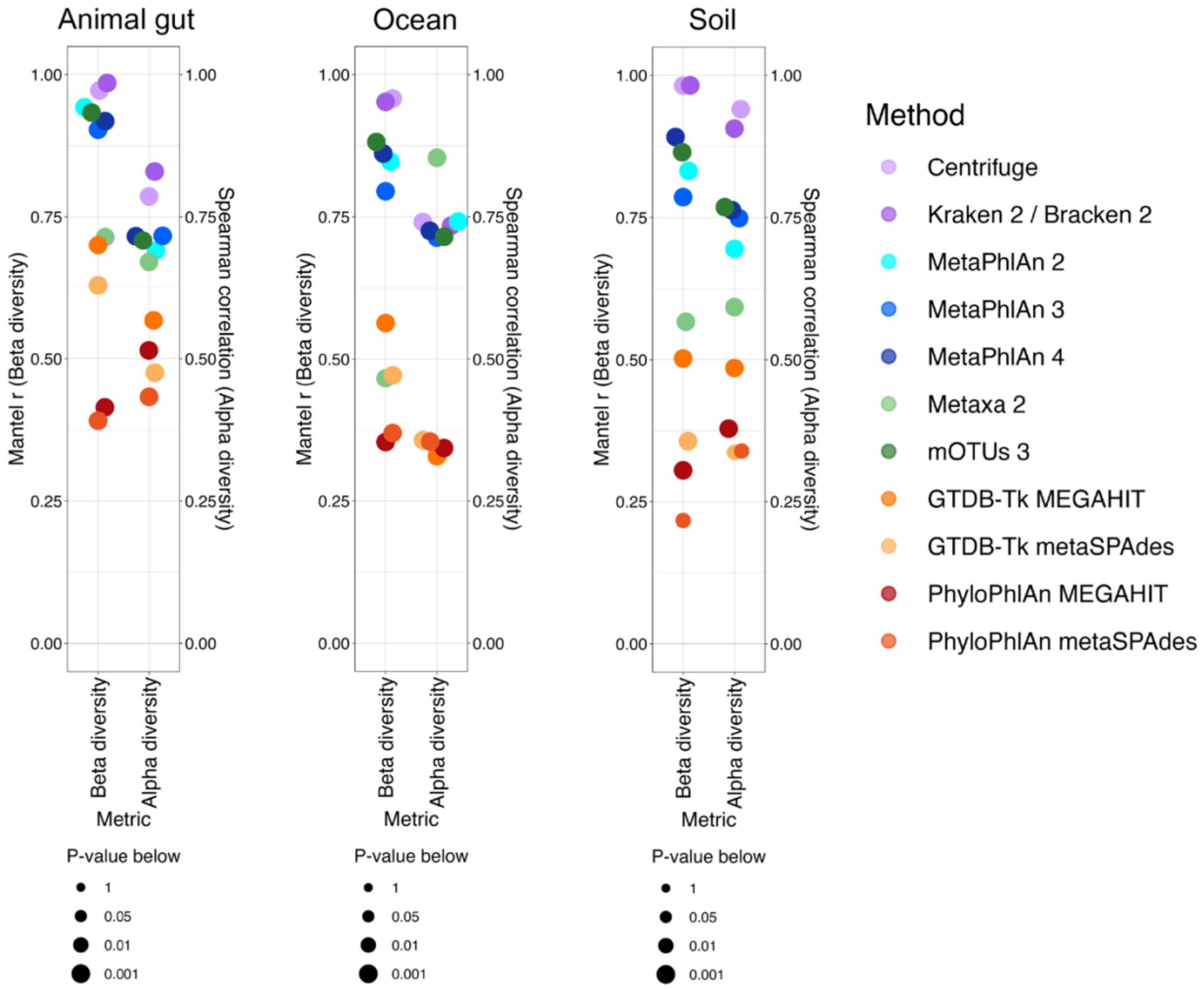

**Supplemental Figure S7: Alpha and beta diversity accuracy.** (Left) Mantel test of the true species Bray-Curtis dissimilarity matrix against the reconstructed Bray-Curtis dissimilarity matrices. Significance was determined from the Mantel test. (Right) Spearman correlation between true and reconstructed inverse Simpson indices at the species level. Significance was determined by permutational testing under a null hypothesis of exchangeability of the diversities assigned to samples. For both metrics, samples' full profiles (NCBI available taxa and NCBI unavailable taxa) were used at the species level.

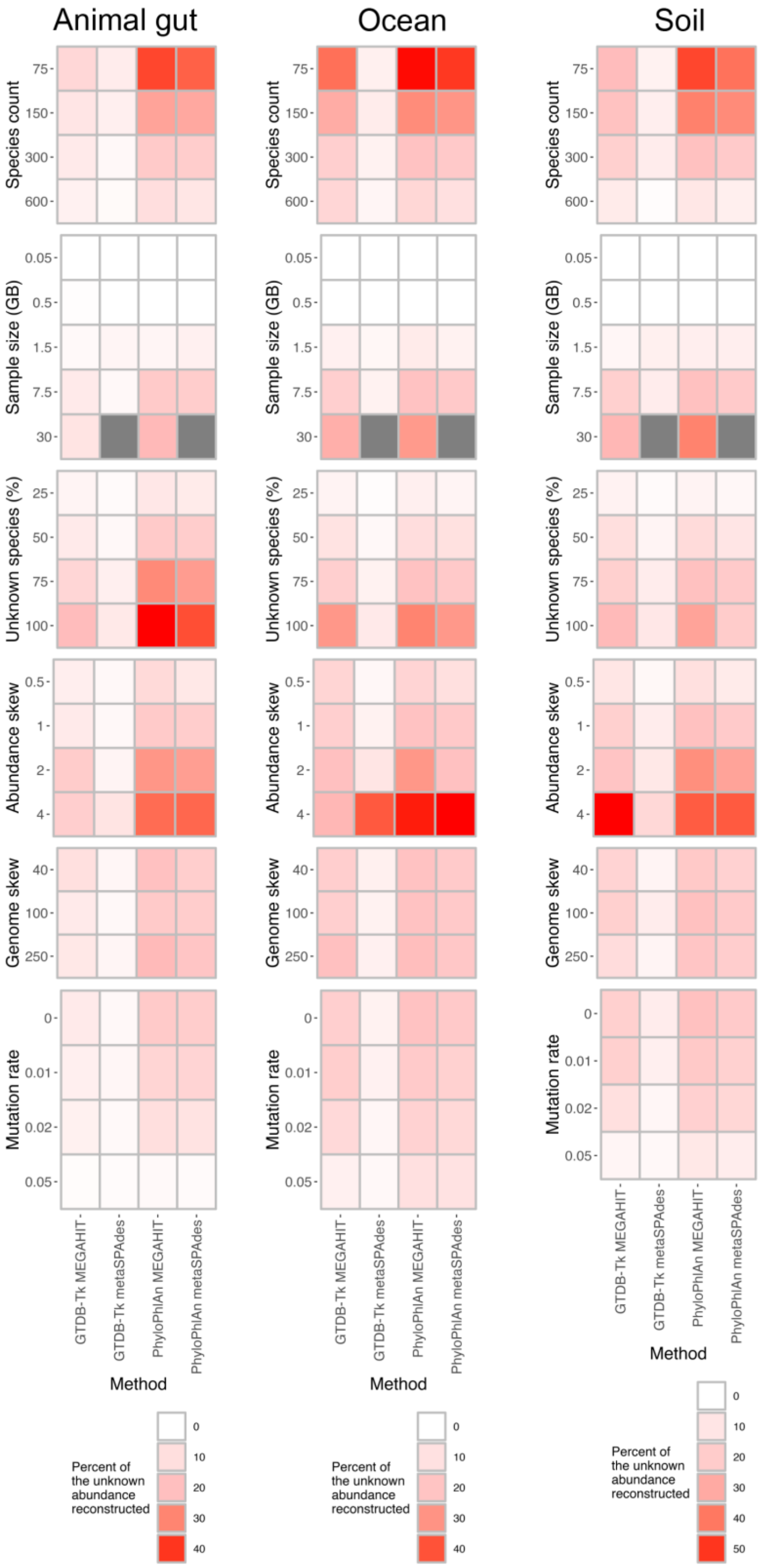

**Supplemental Figure S8: Abundance of uncharacterized species with reconstructed genomes.** Percent abundance of species with genomes of unknown origin successfully reconstructed and assigned a proper taxonomy. Gray boxes indicate that the parameter setting was computationally infeasible to run.

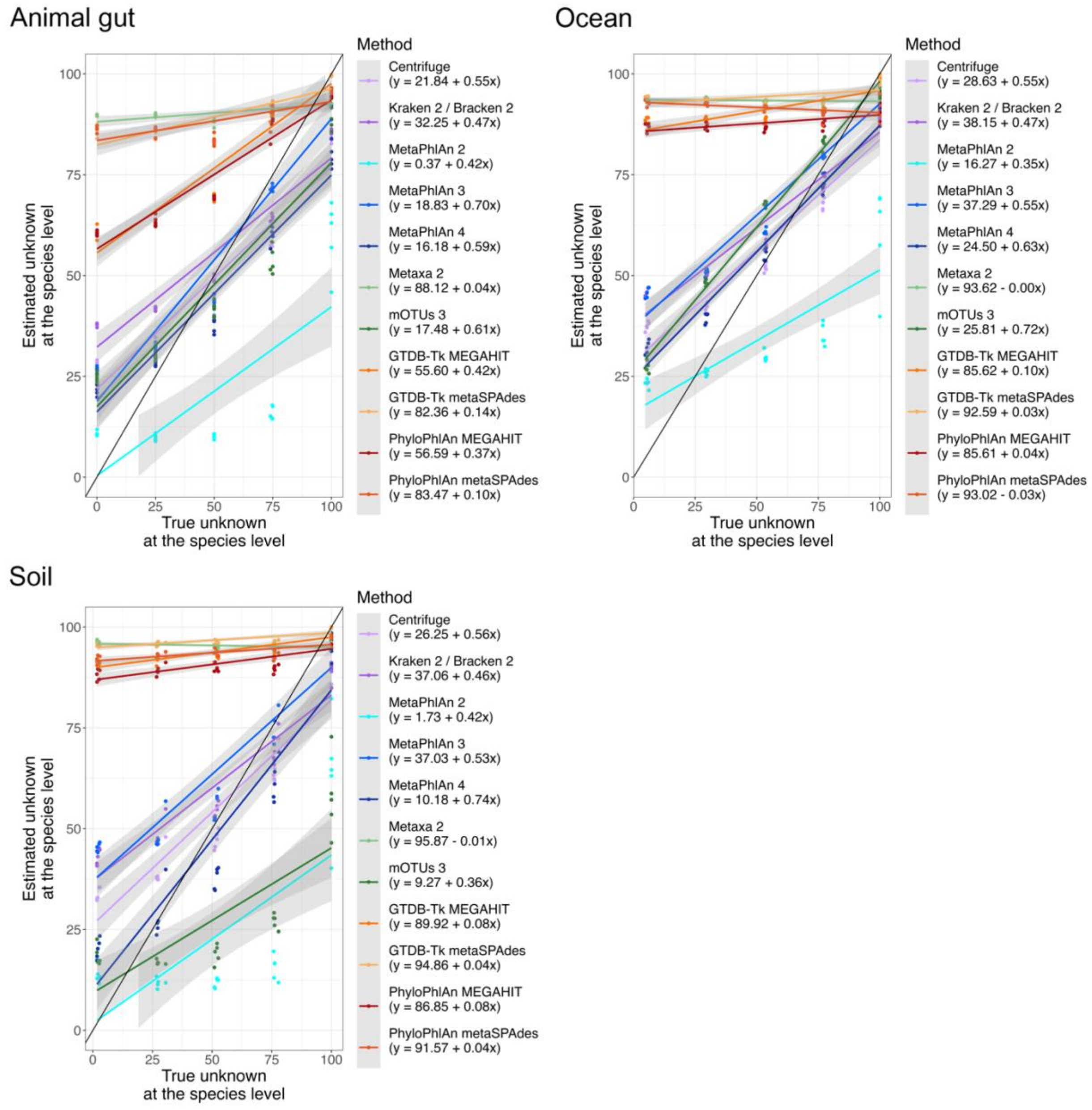

**Supplemental Figure S9: Estimated versus true unknown abundance.** Estimated proportion of the abundance unknown at the species level versus the true abundance of species new to NCBI or from new SGBs.

### The bad: taxonomic profiling methods assign substantially different profiles to the same sample

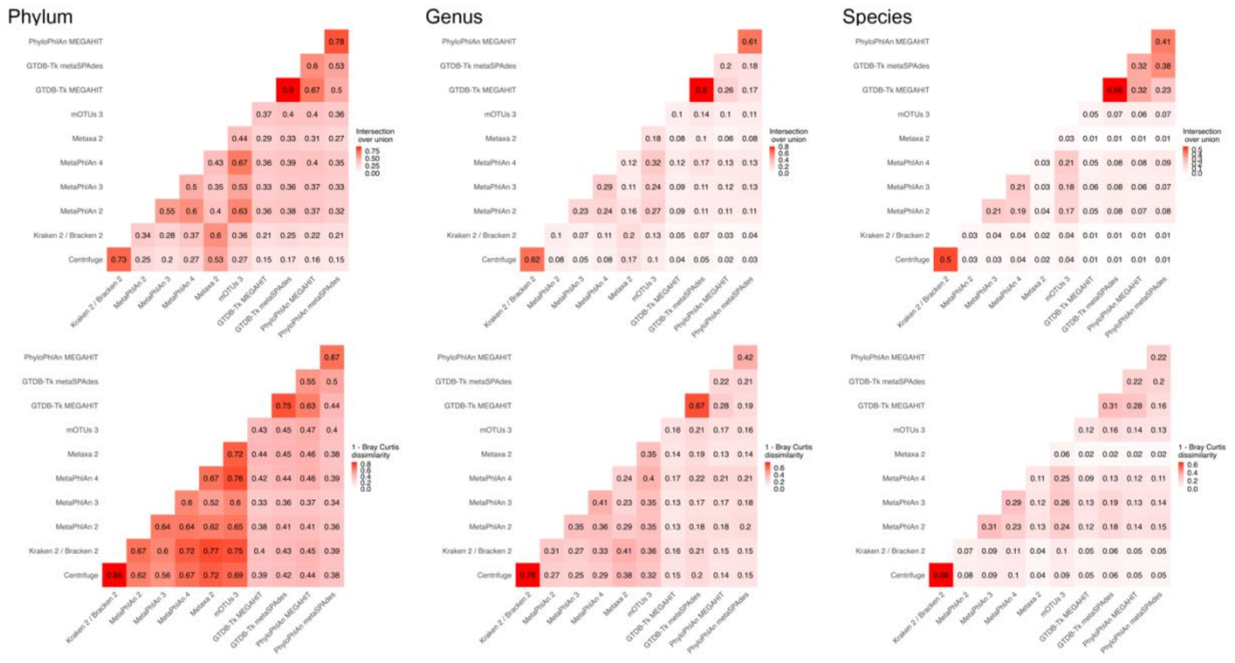

**Supplemental Figure S10: Agreement of methods on real samples.** Average intersection over union and average Bray-Curtis dissimilarity for assignments on the same sample averaged over the non-human datasets.

### The ugly: different methods produce different microbial community structures

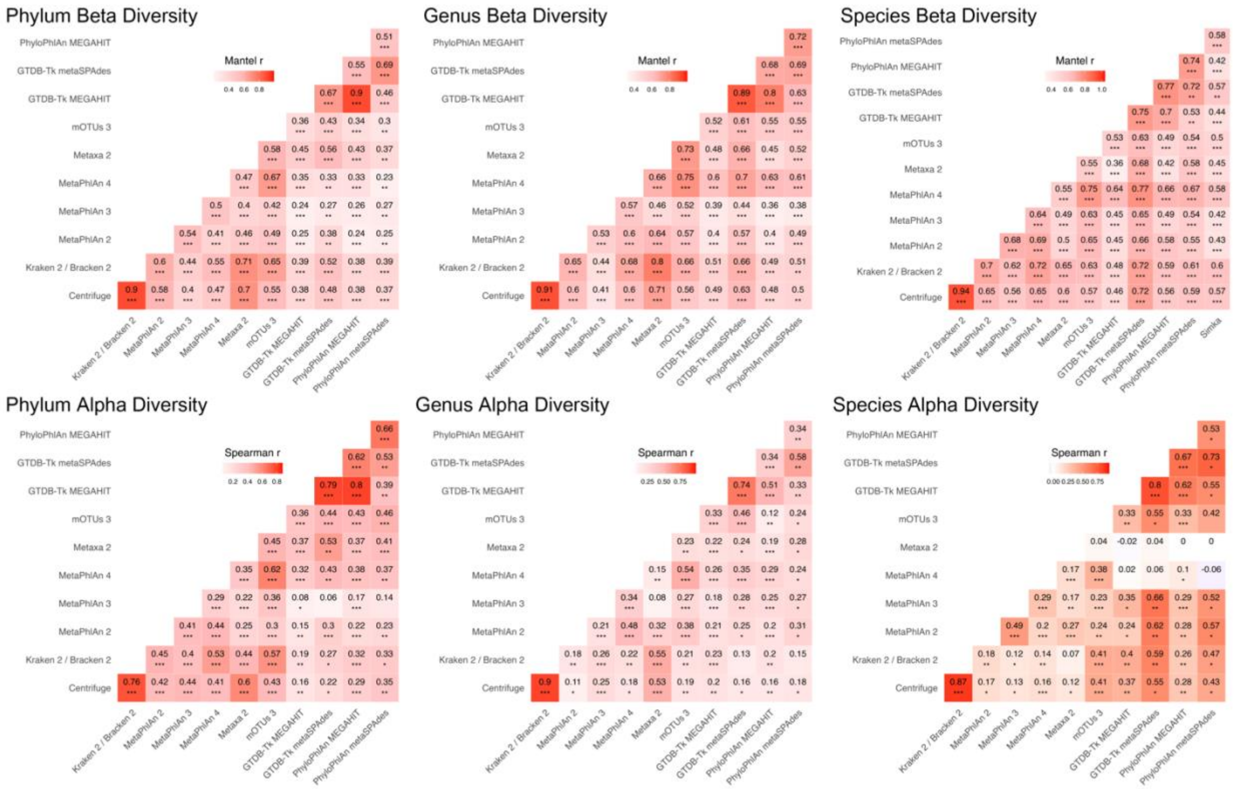

**Supplemental Figure S11: Inferred community structure comparisons.** For beta diversity, a Bray-Curtis dissimilarity matrix for all samples in each dataset was calculated from each method's assignments, and pairs of matrices were compared. Mantel r values are the (unweighted) average of Mantel r values across non-human dataset. Significance is from a one-sided t-test against the null hypothesis that the true average Mantel r across datasets is at most 0. Stars represent significance (p-value <0.05, <0.01, <0.001). Alpha diversities were computed for each sample from each method's profile, and pairs of methods were compared for each dataset. Spearman r values are the (unweighted) average of Spearman r values across non-human datasets. Significance is from a one-sided t-test against the null hypothesis that the true average Spearman r across datasets is at most 0. For both beta and alpha diversity, all taxa assigned to a sample (not just NCBI-available taxa) were used to compute the Bray-Curtis dissimilarity matrix or inverse Simpson diversity.

### The okay: downstream analysis shows moderate agreement on the most significant effects

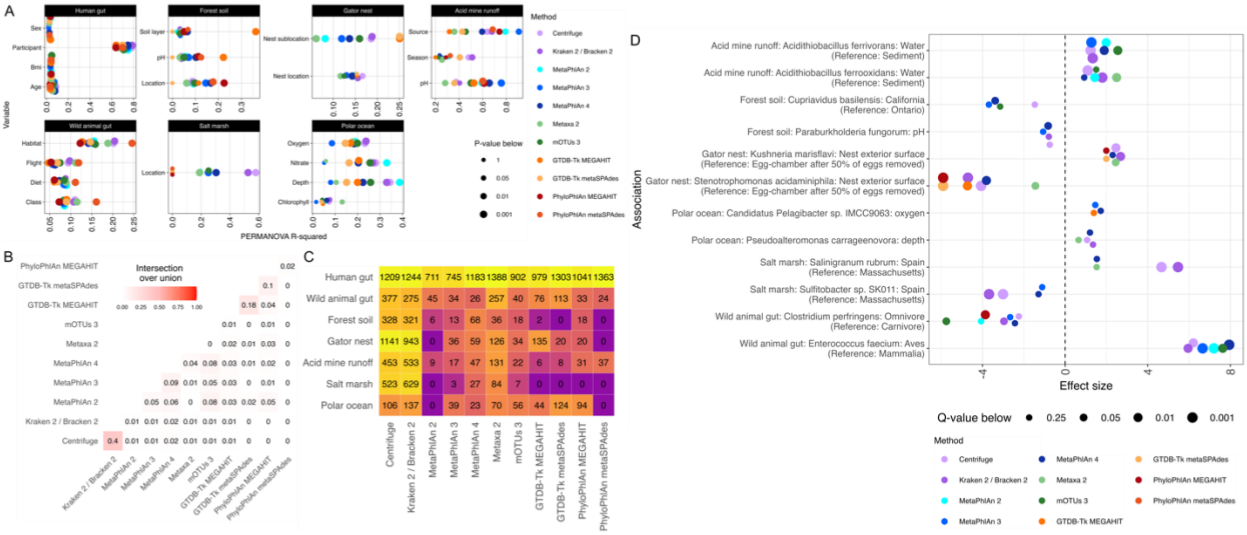

**Supplemental Figure S12: Agreement in identifying important microbial associations. A.** PERMANOVAs of Bray-Curtis dissimilarity for available sample metadata. Human associations were controlled for the subject; all other associations were univariate. **B.** Intersection over union of species-metadata associations determined by MaAsLin 2 averaged (unweighted) over all non-human datasets. **C.** Number of significant species-metadata associations per dataset per method. **D.** For each non-human dataset, the log2 fold changes for the two species-metadata associations identified as significant by the most methods. MaAsLin 2 effect sizes are only shown for a method if the q-value from its profile's association was less than 0.25.

### Discussion

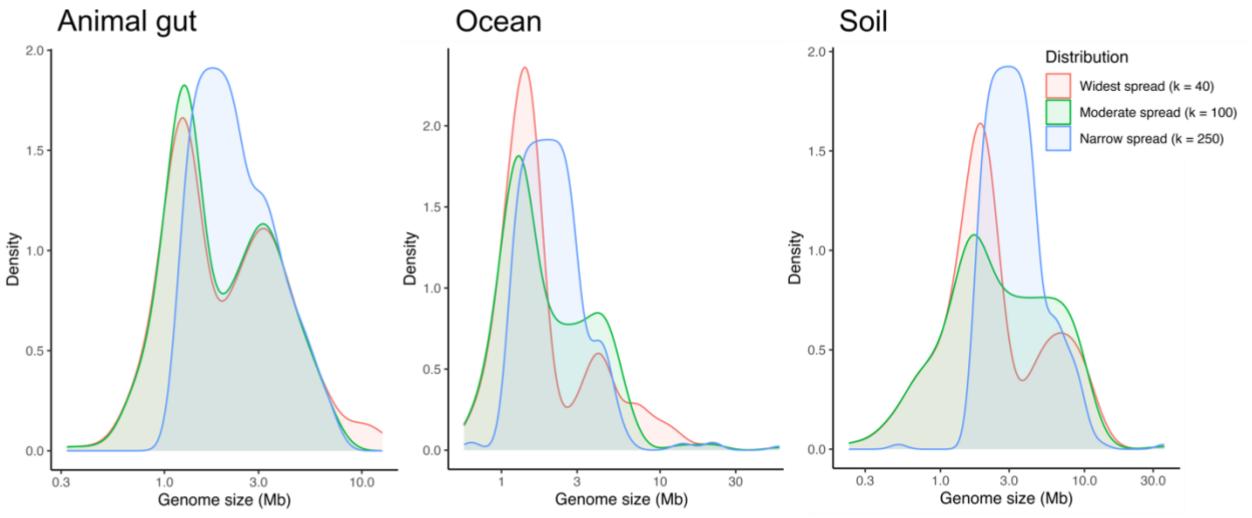

**Supplemental Figure S13: Genome size distribution.** Size distribution for genomes used in each dataset in megabases. One replicate per parameter setting is shown because the distributions are nearly identical within each parameter setting.

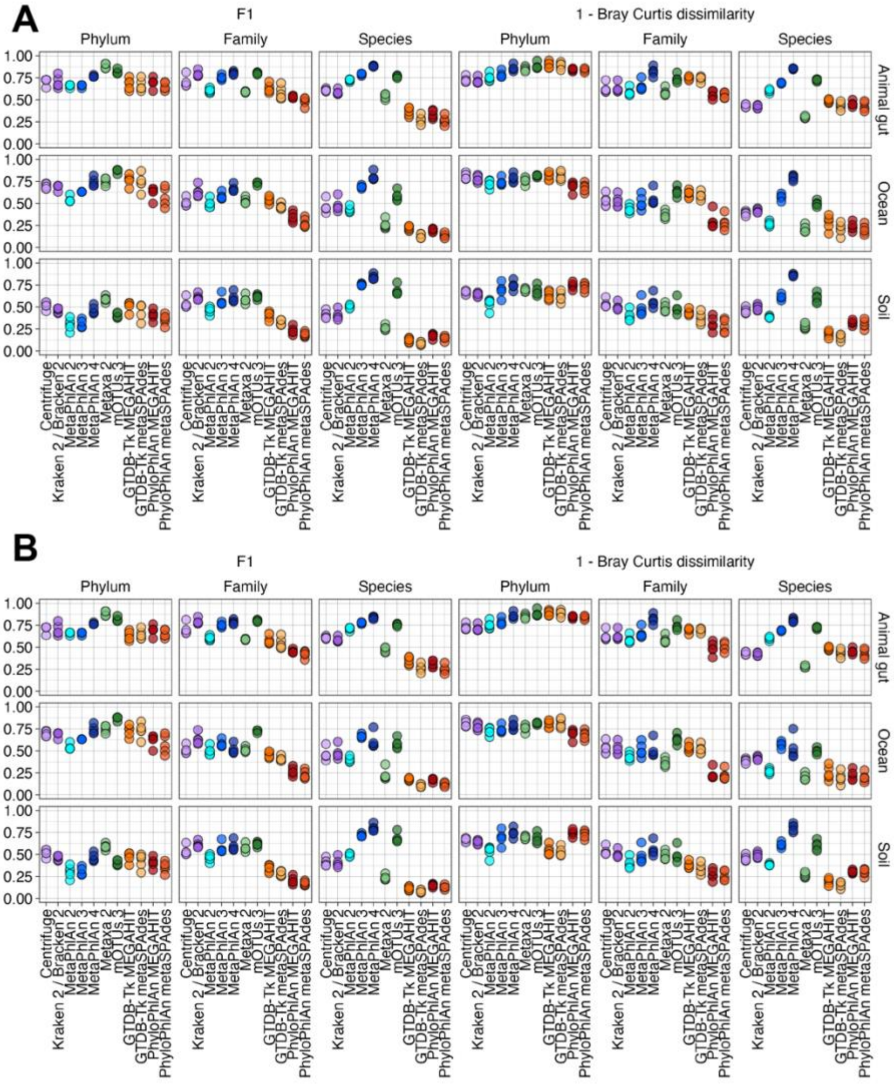

**Supplemental Figure S14: Restricting evaluated taxa to NCBI-available assignments minimally affects accuracy. A.** F1 and Bray-Curtis dissimilarity relative to the true profile for core samples by simulated dataset. Any assigned taxa were retained in the comparison. **B.** Same as A but only assigned taxa with NCBI mappable IDs were retained in the comparison.
